# Differential use of multiple genetic sex determination systems in divergent ecomorphs of an African crater lake cichlid

**DOI:** 10.1101/2021.08.05.455235

**Authors:** Hannah Munby, Tyler Linderoth, Bettina Fischer, Mingliu Du, Grégoire Vernaz, Alexandra M. Tyers, Benjamin P. Ngatunga, Asilatu Shechonge, Hubert Denise, Shane A. McCarthy, Iliana Bista, Eric A. Miska, M. Emília Santos, Martin J. Genner, George F. Turner, Richard Durbin

## Abstract

African cichlid fishes not only exhibit remarkably high rates of speciation but also have some of the fastest evolving sex determination systems in vertebrates. However, little is known empirically in cichlids about the genetic mechanisms generating new sex-determining variants, what forces dictate their fate, the demographic scales at which they evolve, and whether they are related to speciation. To address these questions, we looked for sex-associated loci in full genome data from 647 individuals of *Astatotilapia calliptera* from Lake Masoko, a small isolated crater lake in Tanzania, which contains two distinct ecomorphs of the species. We identified three separate XY systems on recombining chromosomes. Two Y alleles derive from mutations that increase expression of the gonadal soma-derived factor gene (*gsdf*) on chromosome 7; the first is a tandem duplication of the entire gene observed throughout much of the Lake Malawi haplochromine cichlid radiation to which *A. calliptera* belongs, and the second is a 5 kb insertion directly upstream of *gsdf*. Both the latter variant and another 700 bp insertion on chromosome 19 responsible for the third Y allele arose from transposable element insertions. Males belonging to the Masoko deep-water benthic ecomorph are determined exclusively by the *gsdf* duplication, whereas all three Y alleles are used in the Masoko littoral ecomorph, in which they appear to act antagonistically among males with different amounts of benthic admixture. This antagonism in the face of ongoing admixture may be important for sustaining multifactorial sex determination in Lake Masoko. In addition to identifying the molecular basis of three coexisting sex determining alleles, these results demonstrate that genetic interactions between Y alleles and genetic background can potentially affect fitness and adaptive evolution.

## Introduction

Sex, as a means of generating beneficial combinations of alleles, is one of the most effective evolutionary innovations used among eukaryotes to surmount fitness challenges. Many different means of establishing separate sexes have arisen across the tree of life, operating through a combination of genetic and environmental mechanisms (Bachtrog *et al*., 2014; Pennell *et al*., 2018). The continued evolution of new sex determination systems can provide a means to improve fitness via altering sex ratios (Kocher, 2004), resolving sexually antagonistic mutations (van Doorn & Kirkpatrick, 2007; 2010), and avoiding the negative consequences of sex chromosome degeneration (Blaser *et al*., 2013). Given this adaptive role of sex determination, this begs the question of whether it is any coincidence that the fastest reported rates of sex chromosome and heterogamety transitions among vertebrates (El Taher *et al*., 2020) have occurred in East African cichlid fishes, renowned also for their extremely high speciation rates (Brawand *et al*., 2014; Ronco *et al*., 2020). In support of such an association, population genetic models have demonstrated how heterogamety switches arising from a new sex-determining locus coupled with sexual and sex-ratio selection can help generate reproductive isolation in sympatry (Lande *et al*., 2001).

Sex-determination across African cichlid species is largely governed genetically in either a single-locus or polygenic fashion (Ser *et al*., 2010). The loci controlling sex are known to exist both on homomorphic sex chromosomes, for which there is little if any evidence for long range suppression of recombination around the sex-determining alleles (Parnell & Streelman, 2013), and on supernumerary B chromosomes (Clark *et al*., 2017; Clark & Kocher, 2019). Within the Lake Malawi haplochromine cichlid radiation, the characterized sex determining loci are the orange blotch associated ZW locus and an XY locus on chr5 (Roberts *et al*., 2009; Ser *et al*., 2010), two XY loci on chr7 (Albertson, 2002; Parnell & Streelman, 2013; Roberts *et al*., 2009), an XY locus on chr3, and a ZW locus on chr20 (Parnell & Streelman, 2013), using the chromosome numbering established for the *Metriaclima zebra* genome (Conte & Kocher, 2015). In most of these cases, multiple sex determination systems have been observed to act within a single species. Most studies to date have identified sex-associated loci through captive-breeding experiments (e.g. Parnell & Streelman, 2013; Ser *et al*., 2010), which provide only broad genomic resolution, or through GWAS on relatively small sample sizes in wild populations with limited power to detect intraspecific associations (El Taher *et al*., 2020). While these studies point to cichlid sex determination as being highly fluid on the timescale of hundreds of thousands to millions of years, studies on the dynamics within populations would provide the context for examining how recombination, selection, and drift interact with molecular mechanisms to shape the evolution of nascent sex chromosomes (Furman *et al*., 2020). To this end, we sought to understand how sex determination acts in a single population of the eastern happy cichlid *Astatotilapia calliptera*.

*Astatotilapia calliptera* is found both in the shallow margins of Lake Malawi as well as in the surrounding rivers and smaller lakes. Peterson *et al*. (2017) found that the major chr7 XY locus previously identified in Malawi Mbuna cichlids determined sex in a population of *A. calliptera* from Lake Malawi. Despite only mapping the effect to megabase-scale resolution, they postulated that a variant in the gonadal soma-derived factor (*gsdf*) gene on chromosome 7 was responsible for dictating sex given its repeated role in sex determination in other fish species (Einfeldt *et al*., 2021; Jiang *et al*., 2016; Kaneko *et al*., 2015; Myosho *et al*., 2012).

In particular, we studied *A. calliptera* in crater Lake Masoko to the north of Lake Malawi, which is estimated to have formed ~50,000 years ago (Williamson *et al*., 1999). Lake Masoko is only 700 metres in diameter with a shallow littoral margin and walls steeply descending to around 36 m at its deepest point (Turner *et al*., 2019). It is currently a closed system, without surface connections to any other water bodies (Turner *et al*., 2019). With the only other fish being two cichlid species distantly related to *A. calliptera* and one clariid catfish species, the lake provides a relatively simple context for studying the evolutionary genetics of sex determination, speciation and their potential interaction. Genomic evidence suggests that *A. calliptera* colonised the shallow littoral habitat from nearby river systems ~10,000 years ago, and subsequently extended its range into the deeper benthic habitat ~1,000 years ago (Malinsky *et al*., 2015). These shallow littoral and deep benthic populations are phenotypically distinct ecomorphs, with the differences in habitat use coinciding with differences in body shape and jaw morphology. Moreover, the ecomorphs can be distinguished by differences in male breeding colouration, with reproductively active littoral males being typically yellow, and benthic males dark blue. Both ecomorphs are sexually dimorphic, with males generally larger and more brightly coloured than the females, which tend to have a duller, silvery brown colouration.

## Results

We collected whole genome shotgun sequencing data for 548 *Astatotilapia calliptera* from Lake Masoko at a median coverage of 14.5x (range 4.5x - 22x, mean of 12.2x), and combined this with data from 99 previously published samples (Malinsky *et al*., 2015), resulting in whole genome sequence data for 596 male and 51 female fish (Supplementary Table 1). European Nucleotide Archive accessions for the raw Lake Masoko *A. calliptera* sequence data are provided in Supplementary Table 1. Reads were mapped to the high-quality fAstCal1.2 *A. calliptera* reference genome and variants called at 3,328,052 quality-screened single nucleotide polymorphism (SNP) sites (see Methods for details). All commands, code, and links to downloadable source data used to generate the following results and figures can be found at https://github.com/tplinderoth/cichlids/tree/master/Masoko_sex_study.

**Table 1:**
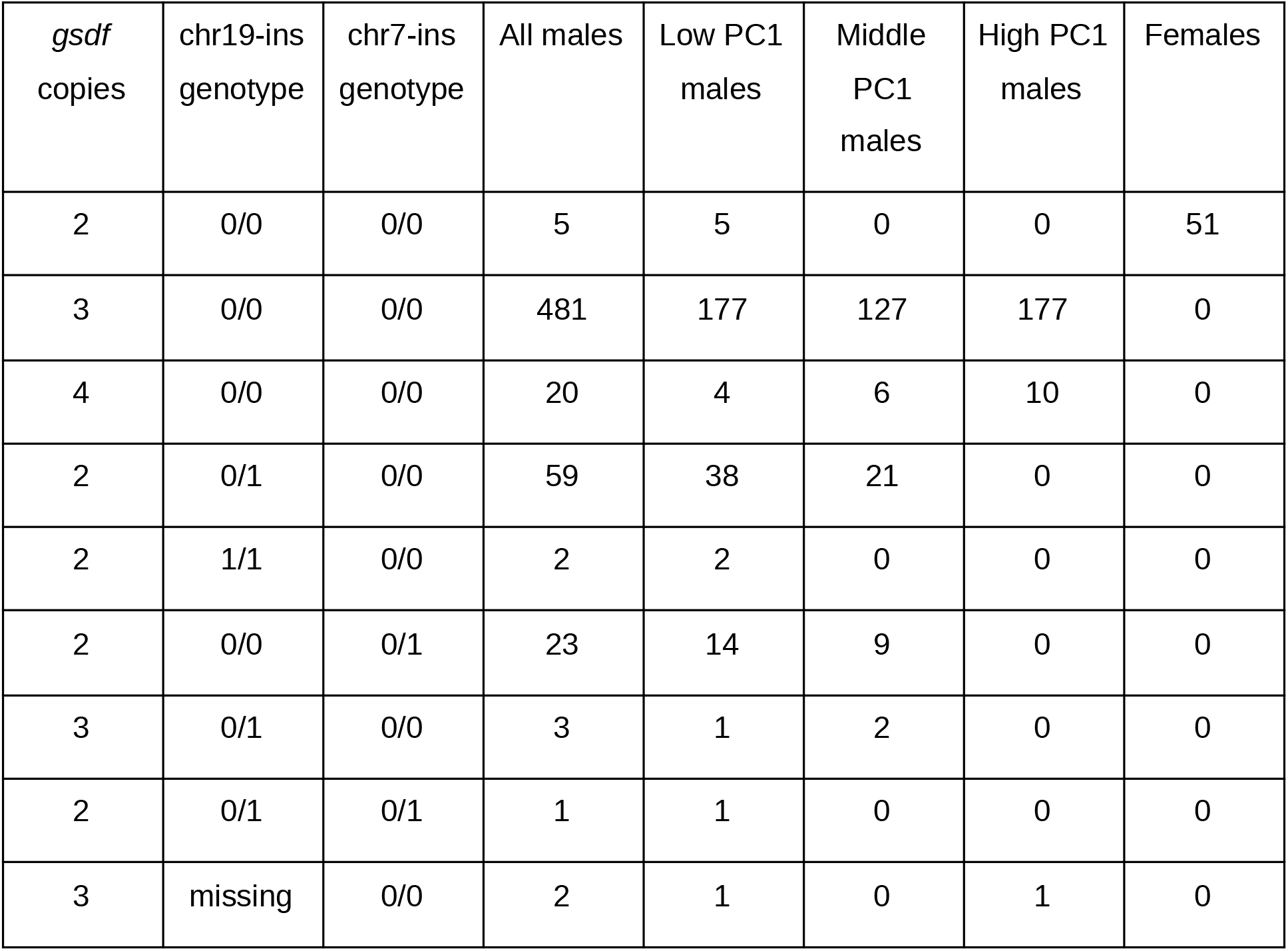
Frequency of sex-determining genotypes in Lake Masoko *Astatotilapia calliptera*. Multilocus genotypes for the sex determining loci are based on the number *gsdf* gene copies an individual carries and their combination of reference (0) and insertion (1) alleles at the loci characterized by the chr19-ins and chr7-ins alleles. Among the 51 females in our sample, 46 were classified as low PC1 and five were middle PC1, none of which carried the *gsdf* duplication nor any of the insertion alleles.

| *gsdf* copies | chr19-ins genotype | chr7-ins genotype | All males | Low PC1 males | Middle PC1 males | High PC1 males | Females |
| --- | --- | --- | --- | --- | --- | --- | --- |
| 2 | 0/0 | 0/0 | 5 | 5 | 0 | 0 | 51 |
| 3 | 0/0 | 0/0 | 481 | 177 | 127 | 177 | 0 |
| 4 | 0/0 | 0/0 | 20 | 4 | 6 | 10 | 0 |
| 2 | 0/1 | 0/0 | 59 | 38 | 21 | 0 | 0 |
| 2 | 1/1 | 0/0 | 2 | 2 | 0 | 0 | 0 |
| 2 | 0/0 | 0/1 | 23 | 14 | 9 | 0 | 0 |
| 3 | 0/1 | 0/0 | 3 | 1 | 2 | 0 | 0 |
| 2 | 0/1 | 0/1 | 1 | 1 | 0 | 0 | 0 |
| 3 | missing | 0/0 | 2 | 1 | 0 | 1 | 0 |

### Multiple Y alleles determine sex in Lake Masoko

We carried out a genome wide association study (GWAS) for sex using a linear mixed model framework (Figure 1a). The most strongly associated SNP is very highly significant (log_10_ p-value = 2.02e-22), and located at position 18,098,212 on chromosome 7 approximately 8 kb downstream of the gene *gsdf*. By considering read depth summed over all fish heterozygous for this SNP, we established that it, and the entire *gsdf* gene, are contained in a 20 kb-long region that exhibits 50% inflated relative coverage in the heterozygotes, suggesting that the associated variant chromosome contains a duplication of this region (Figure 1b). We examined paired end Illumina reads from Masoko *A. calliptera* samples homozygous for the apparent duplication (Supplementary Figure 1a), and long Pacific Biosciences reads from a male fish from a related species (*Tropheops* sp. ‘mauve’) which also shows the inflated coverage pattern (Supplementary Figure 1b), and in both cases confirmed the presence of a tandem duplication spanning coordinates 18,079,155 to 18,100,834 of chr7. We also confirmed the presence of this duplication junction by PCR (Supplementary Figure 1c). Copy number of the duplication is a stronger predictor of sex than the best associated SNP from the GWAS scan (Table 1), suggesting that the duplication itself operates as a Y allele in an XY sex determination system.

**Figure 1:**
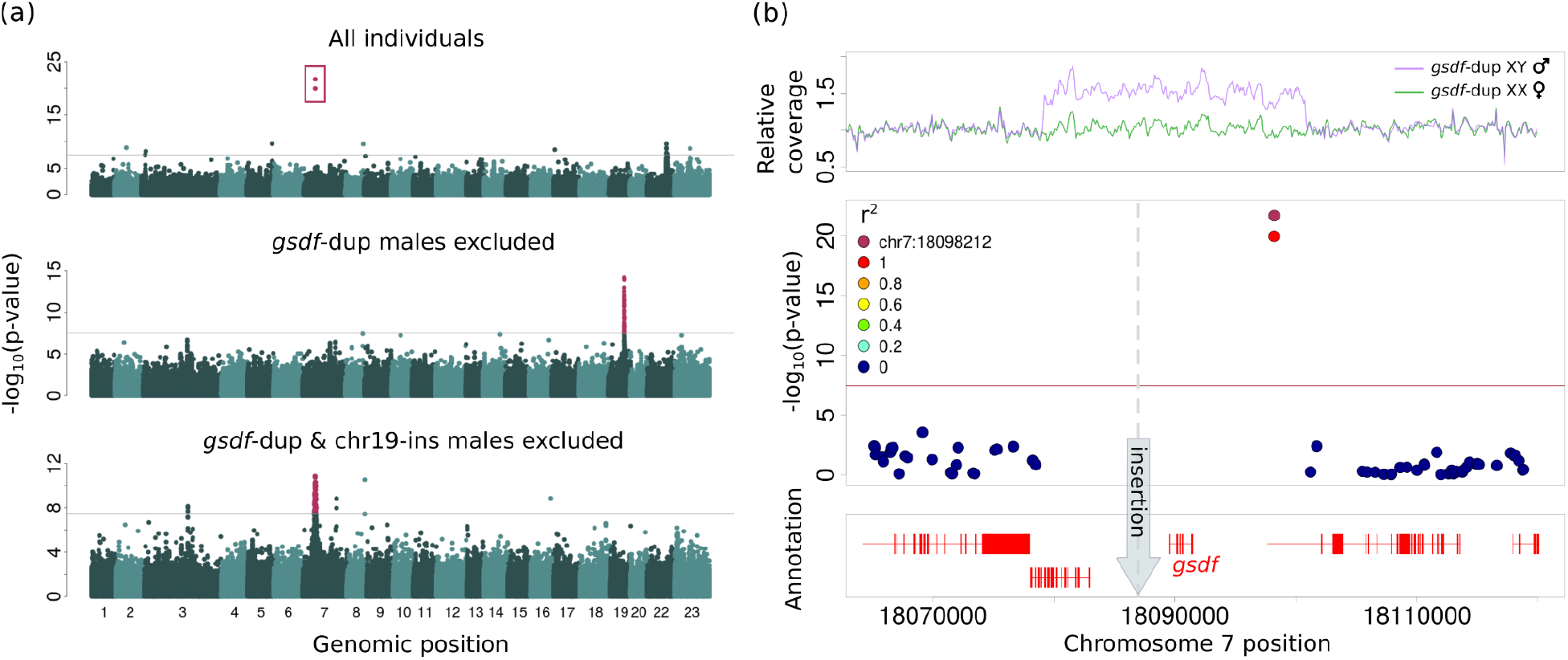
Genome-wide association study for sex. **(a)** P-values for the likelihood ratio test of an association between sex of *Astatotliapia calliptera* from Lake Masoko and their posterior mean genotypes at SNPs across the genome. The panels in order from top to bottom show results from the serial GWAS in which we looked for sex associations using all females and a subset of males not possessing the alternate allele of the single most highly-ranked SNP (or *gsdf*-dup specifically for iterations two and three) from any of the previous GWAS. The grey, horizontal line in each of the Manhattan plots indicates the 0.05 Bonferroni-adjusted significance threshold, correcting for the number of tested SNPs. Significant SNPs tagging sex-determining loci are shown in maroon. **(b)** A zoomed-in view of the region harboring the SNPs most strongly associated with sex on chromosome 7. SNPs are coloured based on their degree of linkage disequilibrium with the most strongly sex-associated SNP tagging the *gsdf* duplication. The top panel shows the average sequencing depth in 100 bp bins of males heterozygous for the *gsdf* duplication compared to females. The sequencing depth of each individual was normalized with respect to their average depth in the non-duplicated flanking regions such that an increase of 0.5x in males compared to females indicates the presence of an extra copy of this locus. The duplication spans the region containing the entire *gsdf* gene and SNPs just downstream of *gsdf* were highly associated with sex in the GWAS run on all males and females. A 5 kb insertion upstream of *gsdf* indicated by the grey arrow characterizes the chr7-ins Y allele, which was in high linkage with the strongly sex-associated chromosome 7 SNPs in the bottom panel of (a).

The duplicated *gsdf* Y allele, which we call *gsdf*-dup, does not determine sex in all males: 90 of the 596 males (15%) are homozygous unduplicated, while 20 (3%) are apparently homozygous duplicated (2x relative sequence depth). To establish whether another locus might control sex in the males lacking *gsdf*-dup, we carried out a second sex GWAS with the 51 females and 90 males without the duplication. This revealed a region on chromosome 19 with multiple SNPs that were highly significant, the highest of which (position 21,581,905, log_10_ p-value = 6.327883e-15) is located 77 bp upstream of the *e2f2* gene (Figure 1a). The inferred ancestral allele at this SNP was found exclusively among males across 59 heterozygotes and 3 homozygotes, suggesting a second XY system (Supplemental Table 2). We inspected the genomic region harboring variants in high linkage disequilibrium (LD) with the SNP to determine whether it was tagging any other variants having an even stronger sex association not detected by the GWAS, which was limited to biallelic SNPs. We discovered one such variant, a 700 bp insertion at position 21,572,413, which is located 1.7 kb upstream of the *id3* gene (Supplementary Figure 2). This male-exclusive insertion, hereafter called chr19-ins, is found in 62 of the 90 males without *gsdf*-dup, of which 60 are heterozygotes and two are homozygotes. There are also three males with *gsdf*-dup that are heterozygous for chr19-ins. The additional sequence inserted in chr19-ins occurs in 37 places across 17 chromosomes and two unplaced scaffolds of the reference genome (blastn evalue = 0, > 96% identity, 100% coverage), and matches an LTR/Unknown family transposable element (blastn evalue = 0, 97% identity, 99% coverage) identified by repeatModeler2. At a more relaxed level of identity this transposable element is found in 126 places spread across all chromosomes and eight scaffolds of the reference genome (blastn evalue = 0, > 92% identity, 100% coverage).

Since there remain 28 males carrying neither *gsdf*-dup nor chr19-ins, we repeated the GWAS procedure a third time, yielding another highly significant region of association on chromosome 7 around *gsdf* (Figure 1a). The most significant individual SNP in this case is approximately 371 kb upstream of *gsdf* (position 17,718,711, log_10_ p-value = 1.386670e-11), with a derived allele exclusively in males; 19 of the 28 males are heterozygous and one is homozygous (Supplemental Table 2). This pattern is consistent with a third Y allele that affects the *gsdf* gene independently of the *gsdf* duplication. Further investigation in the window of elevated LD with this top GWAS SNP revealed a 5 kb insertion at position 18,086,980, hereafter called chr7-ins, located just 2.5 kb upstream of *gsdf*. This insertion is again exclusive to males including all with the chr7:17718711 derived allele as well as three additional males without any previously identified Y allele. Two subregions of the chr7-ins sequence, one 638 bp and the other 510 bp, are respectively found at 19 and 18 places throughout 15 chromosomes and three unplaced scaffolds of the *A. calliptera* reference genome (blastn evalue = 0, >90% identity, 100% coverage). RepeatModeler2 assigns them both to the ends of an unknown repeat family, indicating that the chr7-ins insertion was also introduced by a transposable element. There remain 5 males (0.8% of 596) not carrying any of the three putative Y alleles (*gsdf*-dup, chr19-ins, chr7-ins). These results showing all genotypes are summarized in Table 1. It has been reported that B chromosomes can act dominantly to determine female sex in some rock-dwelling Mbuna Lake Malawi cichlids (Clark *et al*., 2017; 2018; 2019). We therefore examined whether any of our Lake Masoko samples contained excess sequence indicative of B chromosomes, as defined in Clark *et al*. (2018). None of our samples showed any such excess, indicating that B chromosomes do not contribute to sex determination in this system.

### Gsdf is expressed at higher levels in individuals carrying gsdf-affected Y alleles

Comparison of gene expression in the gonads of two adult male and two adult female *A. calliptera* shows seven-fold higher *gsdf* expression in males than in females (Figure 2a), consistent with observations in other fish species of higher levels of *gsdf* in testis than ovary (Zhu et al., 2018). Furthermore, male carriers of *gsd*f-dup and chr7-ins, the latter which could plausibly be in a promoter region of *gsdf* given its upstream proximity, express *gsdf* in non-gonadal tissues (liver, eye, gill and anal fin) at substantially higher levels than males lacking these alleles (Figure 2b & Supplementary Figure 3). Thus, we infer that higher *gsdf* expression resulting from more copies of the actual gene itself or changes to a regulatory element triggers masculinization in Masoko *A. calliptera*. In contrast, the inserted chr19-ins sequence upstream of *id3*, the nearest gene to this insertion, did not show any associated changes in expression. It remains unclear how this variant results in masculinization.

**Figure 2:**
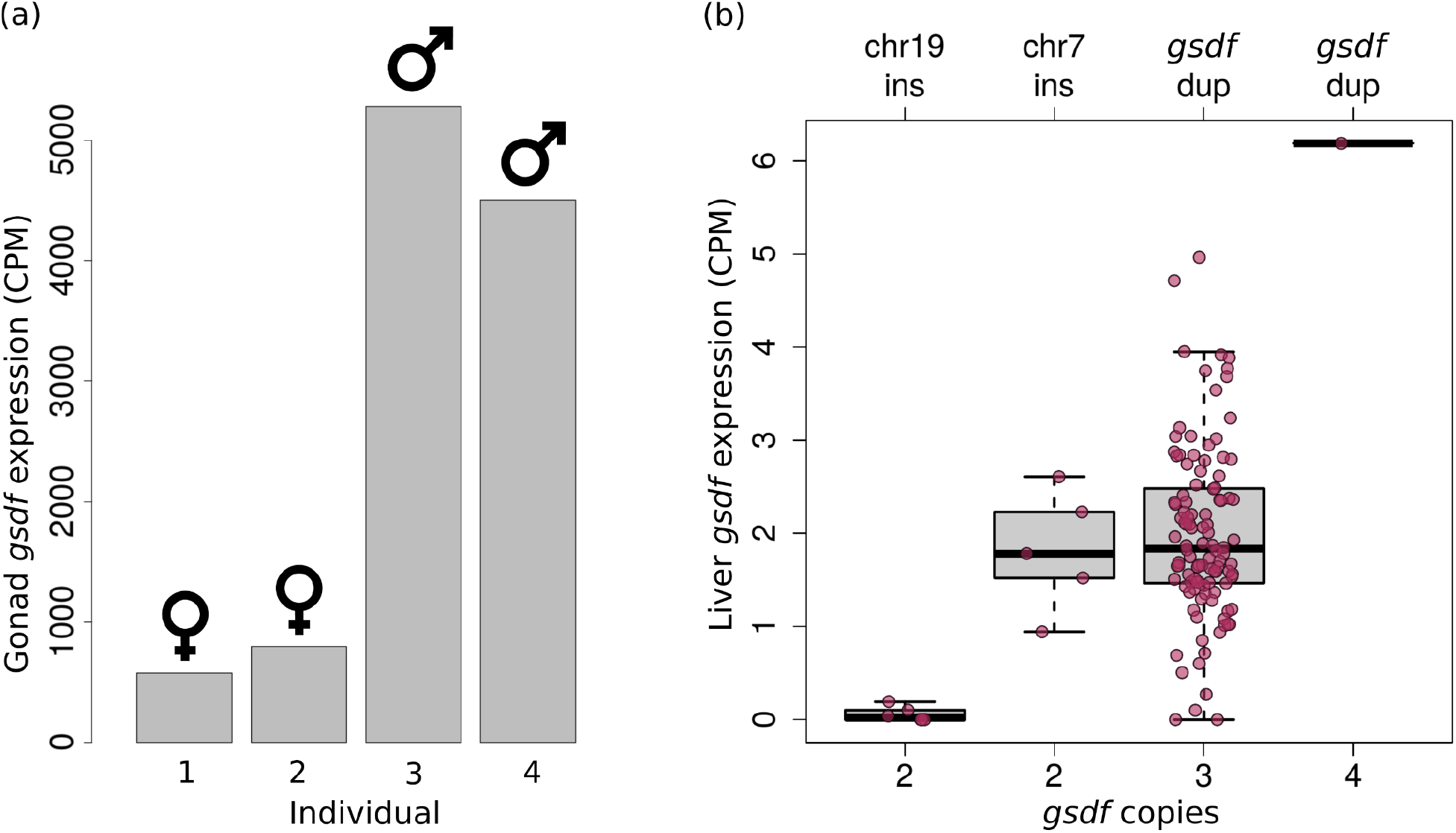
Expression of *gsdf*. **(a)** Expression levels of *gsdf* in the gonads of two male and two female *A. calliptera* reveals approximately seven times higher *gsdf* expression in males. **(b)** Comparison of *gsdf* expression levels in the livers of Masoko male *A. calliptera* heterozygous (three copies) and homozygous (four copies) for the *gsdf* duplication and males lacking the duplication (two copies) but who carry Y alleles generated through insertions on chromosomes 7 and 19. The chromosome 7 insertion (chr7-ins) is directly upstream of *gsdf*, potentially in a regulatory element of this gene. Thus, all males carrying Y alleles resulting from mutations thought to affect *gsdf* express this gene more than other males on average. Gene expression was quantified as counts per million reads (CPM).

### Differential use of Y alleles in Lake Masoko

A principal component analysis (PCA) of the SNP data for the Lake Masoko samples reveals a primary axis of genetic variation distinguishing the benthic from littoral ecomorph (Figure 3a), and this axis is strongly correlated with catch depth (Supplementary Figure 4). There is a tight cluster of samples at high principal component 1 (PC1) corresponding to the benthic ecomorph. For the purposes of this paper we denote fish with PC1 > 0.4 as genetically benthic, and those with PC1 < 0.4 as genetically littoral. The genetically littoral fish are more broadly distributed in the PCA plot, consistent with varying degrees of benthic admixture (Supplementary Figure 5), and for some analyses below we partition them into a “low PC1” subgroup with PC1 < −0.02, and a “middle PC1” group with −0.02 < PC1 < 0.4.

**Figure 3:**
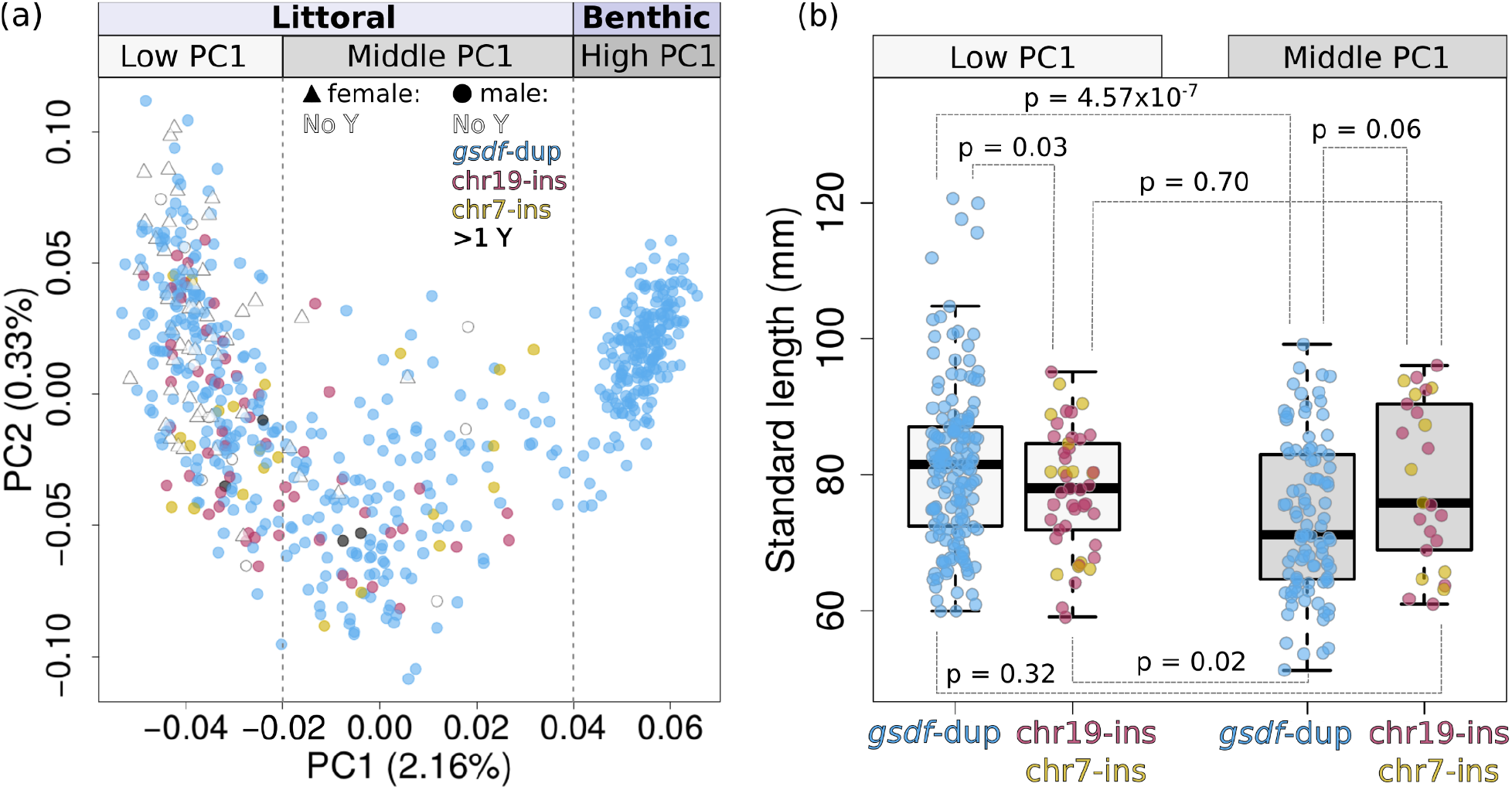
Genetic characterization of Masoko *A. calliptera*. **(a)** The first two components from a principal component analysis of the genome-wide variation among *A. calliptera* from Lake Masoko shows different Y allele usage between fish belonging to distinct genetic clusters. The points represent individuals and their colours denote which of the sex determining alleles identified from the GWAS individuals carry. PC1 separates fish adhering to the benthic ecomorph from littoral morph fish. The dashed grey lines show the demarcations that were used to classify fish as low, middle, and high PC1, which corresponds to their level of benthic ancestry across the genome. **(b)** Comparisons between the standard lengths of littoral males heterozygous for *gsdf*-dup versus males heterozygous for chr19-ins or chr7-ins shows an interaction between Y allele type and benthic admixture levels on body size. Males carrying more than one type of Y allele were excluded. Two-tailed t-tests were used to test for significant differences between the lengths of males characterized by different genetic PC1 background and Y allele combinations (p-values shown).

The genetically benthic fish were almost exclusively found in deep waters (> 20 metres), with just three of 188 individuals at intermediate depth (5-20 metres). The genetically littoral fish were found predominantly at shallow (< 5 metres) and intermediate depths, though there were some littoral fish caught in deep water, with a strong bias for these to be amongst fish with higher PC1 values: in particular, amongst the 289 low PC1 subgroup individuals 138 were caught shallow, 114 at intermediate depth, and 6 deep, while out of the 170 middle PC1 subgroup individuals 25 were caught shallow, 63 at intermediate depth, and 46 deep.

Interestingly, all 188 genetically benthic males carried the *gsdf* duplication compared to 318/408 (78%) of the remaining males (Figure 3a); this deviates significantly from a null hypothesis in which the frequency of males using *gsdf*-dup is independent of PC1 (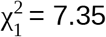, p = 0.007). Correspondingly, the chr19-ins and chr7-ins alleles are only present in the genetically littoral males, at respective frequencies of 8.2% and 2.9%.

### Antagonism between Y alleles and admixture

Fish grow throughout life, and there is evidence that physical size is a correlate of resource holding potential and reproductive success in males of African mouthbrooding cichlids (Hermann et al., 2015; Nelson, 1995; Sefc, 2011) where even a 1 mm size difference can severely impact an individual’s chances of winning bouts of male-male aggression (Turner & Huntingford, 1986). In Lake Malawi haplochromine cichlids specifically, body size is a key predictor of the ability to successfully hold essential breeding territory from which to court females (Markert & Arnegard 2007). Even in the absence of male-male competition, at least in the case of South American convict cichlids, females prefer to mate with larger males (Dechaume-Moncharmont *et al*., 2011), thus there is substantial evidence to suggest that male cichlids may commonly benefit from being larger.

In Lake Masoko, the genetically littoral male fish tend to be smaller as their amount of benthic ancestry increases (Supplementary Figure 6, Supplementary Table 3). This decrease in size with greater benthic admixture is significantly influenced by the type of Y allele that a male carries (ANOVA F = 3.66, p = 0.027, comparing a linear model with interaction between genetic PC1 and Y allele to a model with no interaction term). Chr19-ins males and chr7-ins males are the same size in both low and middle PC1 subgroups (low PC1 two-tailed t = −0.40, p = 0.70, middle PC1 two-tailed t = −0.24, p = 0.81), and together their size remains stable regardless of the level of benthic ancestry (two-tailed t = 0.38, p = 0.7, Figure 3b). In contrast, *gsdf*-dup males with middle PC1 genetic ancestry are significantly smaller than those with low PC1 ancestry (two-tailed t = 5.21, p = 4.57*10^-7^). This size difference for *gsdf*-dup males is so pronounced that while they are significantly larger than males using the other two Y alleles on the low PC1 background (two-tailed t = 2.24, p = 0.03) they tend to be smaller in an intermediate PC1 background. In contrast, the *gsdf*-dup genetically benthic (high PC1) males do not suffer from the size deficit seen in *gsdf*-dup middle PC1 males (Supplementary Figure 7a). Males homozygous for *gsdf*-dup are on average 81 mm long, which is no different than heterozygotes (two-tailed t = −0.48, p = 0.64), and so by this proxy are equally fit.

Because PC1, which reflects benthic genetic content, is correlated with fish capture depth, we examined whether there could be an interaction between environment and genotype contributing to these size differences. Interestingly, while the *gsdf*-dup males with middle PC1 ancestry are smaller at all catch depths, chr19-ins and chr7-ins males with middle PC1 backgrounds are noticeably larger at depths greater than five metres (Supplementary Figure 7a). This larger size of the deeper-caught chr19-ins and chr7-ins middle PC1 males is counteracted by their shallow-caught counterparts tending to be the overall smallest, contributing to these males appearing similar in size across genetic backgrounds when not accounting for depth. Despite numbers of some categories being low, this three-way interaction between the depth at which fish are caught, Y allele type, and level of benthic ancestry, is borderline significant in its ability to predict fish length (ANOVA F = 3.02, p = 0.05), suggesting that depth is relevant in contextualizing how different genetic combinations relate to body size, and therefore fitness.

If the low PC1 and middle PC1 fish were sufficiently separated from each other genetically, these differences in size would be expected to lead to differences in the fraction of littoral males carrying the rarer insertion alleles at greater depth or PC1 values. However, a three-way interaction between PC1 (restricted to low and middle PC1), catch-depth, and Y allele type is not significant in modeling the frequency of males (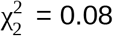, p = 0.96), nor are interactions between Y allele type and depth or PC1 (Wald test z = −0.85 to 1.16, all p-values > 0.25 in the homogeneous association model of male frequency, which includes all pairwise interactions between depth, Y allele and PC1) (Supplementary Figure 7b). Indeed, pooled across depths, *gsdf*-dup males are 3.5x more common than males carrying either of the other two Y alleles among fish with low PC1 genetic backgrounds and 3.9x more common among middle PC1 males (difference not significant, Fisher’s exact test p = 0.45).

Although the results of the last paragraph fail to provide direct evidence of a selective benefit for the Y insertion alleles at deeper depths or highly admixed genetic backgrounds in terms of allele frequency differences, it is noteworthy that elevated linkage disequilibrium (LD) extends for hundreds to thousands of kilobases from the strongest sex-associated GWAS SNPs tagging chr19-ins and chr7-ins (Supplementary Figure 2). To quantify this extent of LD we measured the mean squared physical distance between the chr19-ins and chr7-ins tagging SNPs and other SNPs that were within a megabase and in strong LD (r^2^ > 0.5) with these focal SNPs; these values are in the 81st and 87th percentiles respectively compared to other randomly-sampled focal SNPs across the genome with the same allele frequencies. This is consistent with long-range LD generated by recent positive selection, suggesting that either the sex-determining variants or another locus that they are physically linked to could be the target of selection.

### Distribution of sex-determining alleles across the Lake Malawi cichlid radiation

We next investigated the presence of these Y alleles in other species from the Malawi radiation for which we have sequenced samples. The *gsdf* duplication is seen in 95 additional species, suggesting that it is old and may correspond to the major male-determining allele in the chr7 XY system observed to act previously in multiple Lake Malawi cichlid species (Parnell & Streelman, 2013; Ser *et al*., 2010) (Supplementary Table 5). However, its use in sex determination appears to be quite dynamic; for example, it was not seen in the entire sample of 32 *A. calliptera* males from crater lake Itamba near to Lake Masoko (Figure 4a), and it has been lost or gained multiple times within the *Maylandia* genus (Figure 4b).

**Figure 4:**
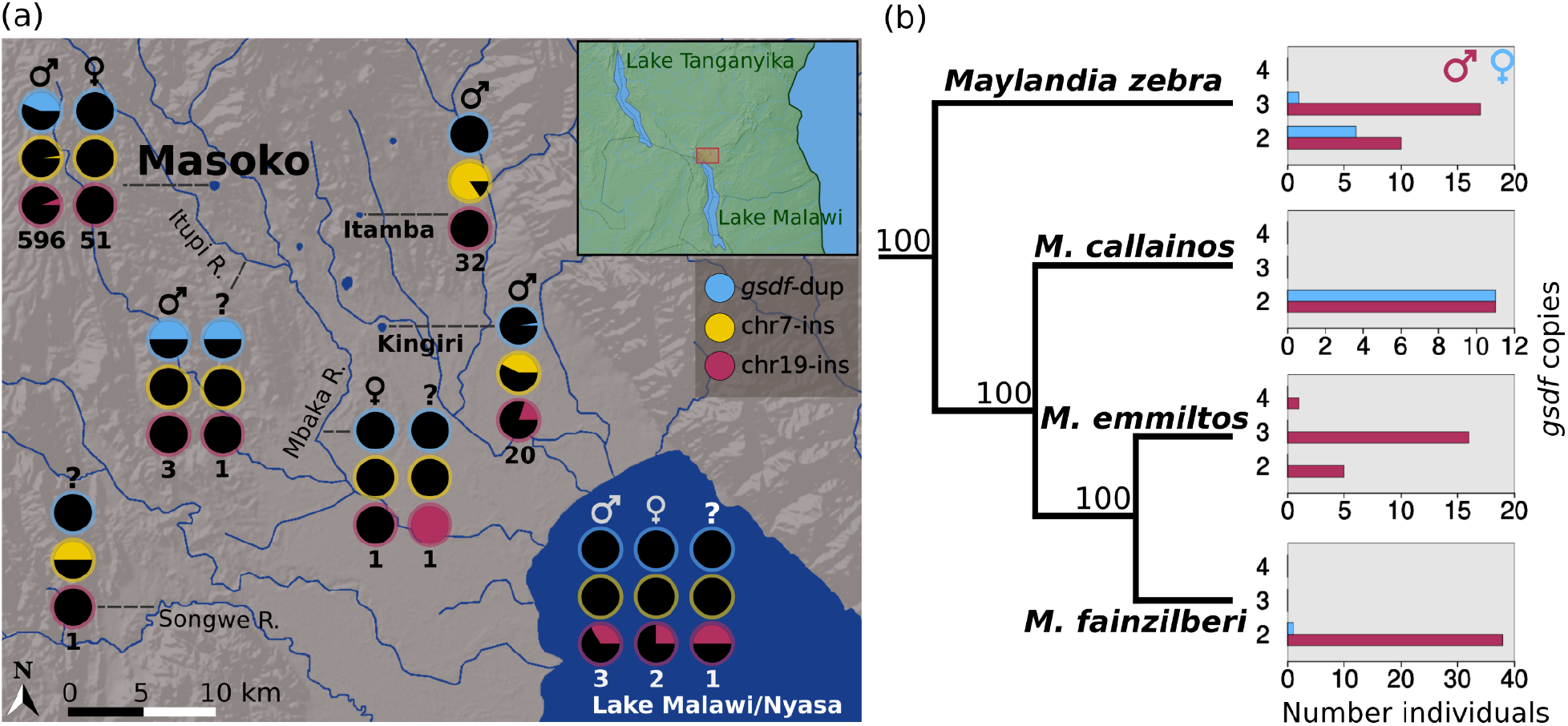
Geographic and taxonomic distribution of Y alleles. **(a)** The frequency of the *gsdf*-dup, chr7-ins, and chr19-ins alleles among *A. calliptera* males, females, and individuals of unknown sex sampled from lakes and rivers throughout Tanzania and Malawi suggests varied usage of these alleles as sex determiners. The sample sizes for each sex and locality are indicated under pie charts of allele frequencies. **(b)** The frequency of male (blue) and female (maroon) individuals from four *Maylandia* species that are either heterozygous (three copy), homozygous (four copy), or lacking (two copy) the duplicated *gsdf* allele exemplifies the dynamic role of *gsdf*-dup in sex determination across the Malawi cichlid radiation. The presence of the *gsdf* duplication in relation to the neighbor-joining species tree, rooted using the distantly-related outgroup *Rhamphochromis longiceps*, suggests that the *gsdf* duplication has been lost or gained at least twice during the diversification of the *Maylandia* lineage. Additionally, the *gsdf* duplication is found in both sexes of *M. zebra*, although at significantly different frequencies (Fisher’s exact test p = 0.035), consistent with it playing a role in sex determination in this population.

Among our specimens, the chr19-ins allele is exclusive to *A. calliptera*, and is geographically widespread, occurring in populations from another Tanzanian crater lake, Kingiri (Figure 4a), as well three other lakes, and five rivers (Supplementary Table 5) that span an area extending south and north of Lake Malawi. Among the 20 non-Masoko chr19-ins carriers for which we have sex information, 18 were chr19-ins heterozygote males from the Bua River and lakes Kingiri, Malombe, Chilwa, and Malawi, and two were heterozygote females from the Salima population of Lake Malawi and the Ruvuma River.

The chr7-ins allele occurs in other lake and riverine populations of *A. calliptera* mostly from the regions surrounding northern Lake Malawi except for one southern Lake Malawi population (Southwest Arm). Among 20 Lake Kingiri males 55% are heterozygous for chr7-ins and 15% are homozygous, while in 32 Lake Itamba males 31% are heterozygous and 69% are homozygous (Figure 4a and Supplementary Table 5). The high frequency of chr7-ins homozygotes, particularly in Itamba, suggests that this variant is either not sex determining or is being epistatically masked by a feminizing allele in these populations. We also detected the chr7-ins variant in seven species from the genus *Tropheops* and two *Pseudotropheus* species (Supplementary Table 6). Both genera are endemic to Lake Malawi and belong to the Mbuna clade that is phylogenetically close to *A. calliptera* (Malinsky *et al*., 2018). Small sample sizes of both males and females for these species and the coincidence of both the *gsdf* duplication and chr7-ins make it difficult to confidently discern whether chr7-ins could be involved in sex determination, although there is an indication in some cases. For instance, there is one *Tropheops gracilior* male without *gsdf*-dup that is heterozygous for chr7-ins while the single female from this species does not carry either of these putative Y alleles. Similarly, in *Tropheops* sp. ‘chilumba’ and *Tropheops* sp. ‘mauve’ there are males heterozygous for chr7-ins without a duplicated *gsdf*, however there are no females for comparison. Such a male is also found from *Tropheops* sp. ‘rust’ but in this species, and *Tropheops* sp. ‘white dorsal’, females occur that carry both *gsdf*-dup and chr7-ins. While sexing errors could be responsible, a potentially more plausible explanation is the presence in *Tropheops* of a dominant female-determining variant at another locus, given that females with one of or both chr7-ins and *gsdf*-dup are observed multiple times. Of the two *Pseudotropheus* species positive for chr7-ins, only one, *Pseudotropheus fuscus*, had sexed individuals; 2/2 males are heterozygous for chr7-ins and have an unduplicated *gsdf*, while the only female lacks both *gsdf*-dup and chr7-ins, which is consistent with chr7-ins being male-determining.

## Discussion

Our genome-wide survey for genetic associations with sex revealed that there are three putative XY determination systems segregating within a single natural population of *Astatotilapia calliptera* from the crater lake Masoko. Among these, two are associated with *gsdf* on chromosome 7: the duplication present in 85% of males, which is the primary mechanism, and an upstream insertion present in 4% of males. The third Y allele is characterized by an insertion on chromosome 19 in 11% of males. These systems are used differentially between the divergent ecomorphs in the lake, with the deep-water benthic morph only using the duplication, while littoral fish use all three systems.

Although use of multiple sex determination systems might seem likely to create sex-ratio biases, multiple Y alleles can coexist without problem in a population, with each male just carrying one of them, and females carrying none of them; Mendelian segregation in the offspring then gives 50% males with the paternal Y and 50% females. Indeed, we saw no females with any of the Y alleles. However in our larger set of males we did detect some that carried two Y alleles, including males homozygous for the *gsdf* duplication and others with two different Y alleles, suggesting that there are some females carrying Y alleles present in the broader population. A possible explanation for this is that a dominant ZW system may also be present at low frequency, in which a dominant feminizing W allele acts epistatically to any of the Y alleles, as seen in some other Lake Malawi cichlid species (Parnell & Streelman, 2013; Ser *et al*., 2010). We did not detect such a W allele in our association scans, possibly because the number of females in our data set did not give sufficient power to detect it at the frequency which would explain our observations. Alternatively, there could be incomplete penetrance of the duplication allele, or genetically male fish could rarely undergo environmentally-induced sex reversal, which has been documented in more taxonomically distant cichlids (Baroiller *et al*., 1995).

Complete genomic sequencing of many wild individuals enabled us to identify the likely causal genetic mechanisms creating new Y alleles and corroborate the suspicion by Peterson *et al.* (2017) that *gsdf* is a sex determination locus in *A. calliptera*. Our findings indicate that the tandem duplication of *gsdf* and the proximal upstream insertion both boost *gsdf* expression, consistent with leading to masculinization as shown in *Oryzias* (Myosho *et al*., 2012). Upregulated *gsdf* expression appears to be generally important for testicular development in fish (Matsuda & Sakaizumi, 2016) and *gsdf* has been reported as a sex determiner in multiple fish species (Einfeldt *et al*., 2021; Jiang *et al*., 2016; Kaneko *et al*., 2015; Myosho *et al*., 2012). Recycling of this gene for sex determination through repeated distinct mutations is evidence for evolutionary conservation of the genetic pathways controlling sex even as the specific sex determining alleles turn over (see Bachtrog *et al.* 2014 and Vicoso 2019 for discussion on this topic). The second gene we identified, *id3*, has not previously been directly associated with sex determination, and while we believe we have identified the responsible mutation we cannot be certain of the affected gene.

The genetic mechanisms generating the Masoko Y alleles parallel those involved in the origin of the *dmy*/*dmrt1bY* male determining gene in *Oryzias latipes*, which arose from a duplication of *dmrt1*. Two transposable elements (TEs) introduced transcription factor binding sites upstream of the *dmrt1b* paralog, which altered its expression leading to it becoming the master sex-determining gene (Herpin *et al*., 2010; Schartl *et al*., 2018). Similarly, both the chr19-ins and chr7-ins Y alleles were created by TE insertions directly upstream of the *id3* and *gsdf* genes respectively, offering support for the notion that TEs may play a potent role in rewiring the expression of genes to function as sex determiners (Dechaud *et al*., 2019).

Usage partitioning among three different Y alleles within a single, isolated population provides a striking example of how dynamic sex determination is in African cichlids. This complements recent work showing that across the Lake Tanganyika cichlid radiation sex systems turn over at a higher rate than previously established for vertebrates (El Taher *et al*., 2020). Previous studies showed that multiple sex determination systems can segregate within captive families involving crosses between Lake Malawi species (Parnell & Streelman, 2013; Ser *et al*., 2010), but did not characterize their distributions within natural populations. Our results from Lake Masoko allow us to explore how multiple co-occurring sex systems segregate in the wild, and their relationship to subpopulation structure.

All of the variants that we identified for controlling sex also exist outside of Lake Masoko. The presence of *gsdf*-dup across all major clades of the Lake Malawi radiation, except for *Diplotaxodon* and *Rhamphochromis*, suggests that it either predated the radiation or arose early in it. Despite this, the *gsdf* duplication has not fixed, instead showing evidence of gains and loss at fine taxonomic scales within genera and even species. In contrast, chr19-ins and chr7-ins are both far more taxonomically constrained, with chr19-ins exclusive to *A. calliptera*, despite being widespread geographically. This suggests that these variants, although at low frequency, are also old and in the case of chr7-ins could have been introduced into *Tropheops* and *Pseudotropheus* through introgression. Another possibility is that chr7-ins, seen in 9/67 (~13%) of the uniquely-classified Mbuna species (2/13 genera) in our dataset, could have arisen in a common ancestor of *A. calliptera* and Mbuna and remained as a minor sex-determining player in comparison to *gsdf*-dup, which we detected in ~75% of the Mbuna species (11/14 genera).

This scenario would suggest that *gsdf*-dup may be selectively advantageous over chr7-ins in most circumstances, while there are some conditions that favour chr7-ins. A common feature of all of the Y alleles we identified is that outside of Masoko they do not always appear to determine sex, suggesting that multifactorial sex determination is common and highly variable with respect to which alleles serve as the major sex determiners, even in closely related species. Having identified some of the precise variants influencing sex differentially across the radiation enables future studies into the evolutionary factors supporting their turnover at a variety of evolutionary scales.

Our results raise the question of which eco-evolutionary contexts promote the invasion and eventual maintenance or loss of new sex determining variants. Theorized evolutionary mechanisms contributing to sex system turnover include resolving sexually antagonistic traits (van Doorn & Kirkpatrick, 2007), escape from deleterious mutational load (Blaser *et al*., 2013), selection on sex ratios (Eshel, 1975), genetic drift (Saunders *et al*., 2018), and transmission distortion (Clark & Kocher, 2019; Werren & Beukeboom, 1998). In considering how our findings align with such models it is important to recognize that we are only observing a snapshot of whatever dynamics may be occurring in Masoko, rather than seeing the evolutionary trajectories of Y allele usage.

Under the classic model of sexually antagonistic selection (van Doorn & Kirkpatrick, 2007), autosomal alleles with differential fitness effects between sexes gain an advantage if they become linked to a new sex determination locus, thus coupling the male-benefiting allele with males and vice versa. The resulting linkage disequilibrium can be reinforced in the long term through reduced recombination in the region containing the sex-determining and sexually antagonistic loci. When multiple sex loci co-occur in a population as in our case, the Y allele conferring the greatest fitness advantage to males will spread.

We found evidence of an antagonistic relationship in terms of body size between the different Y alleles and genetic PC1 in littoral males. In cichlids, larger size confers higher fitness to males by providing them with an advantage in defending spawning sites and procuring access to reproductively active females (Hermann *et al*., 2015). In the shallow waters where spawning littoral fish have been observed, the frequencies of males characterized by different combinations of Y alleles and levels of benthic ancestry correlate well with their average size: *gsdf*-dup males with low benthic ancestry (low PC1) are largest and most common compared to males that either carry the chr19-ins or chr7-ins Y alleles or have more benthic ancestry (middle PC1). This suggests that in shallow water among males with low levels of benthic ancestry, *gsdf*-dup males have a fitness advantage over males that carry the rarer Y alleles. This size advantage disappears however in fish with an increased benthic ancestry component, with middle PC1 *gsdf*-dup males being smaller by nearly 8 mm on average. Furthermore, in waters deeper than five metres, among the fish with middle PC1 ancestry, chr19 and chr7 insertion males actually gain a size advantage over *gsdf*-dup males. These size differences are all greater than the level known to be sufficient for preventing smaller males of another African cichlid species from being able to effectively compete for territories (Turner & Huntingford, 1986). In *A. calliptera* specifically, body size has been shown to significantly influence male-male aggression, presumably because it signals the resource holding potential of competing males (Theis *et al.,* 2015). Therefore, we suggest that the insertion Y alleles may be maintained in the population by a relative advantage under these depth and genetic background conditions, while there is sufficient genetic mixing between the low and middle PC1 subgroups of littorals to prevent establishment of significant allele frequency differences.

We suggest two possible reasons, not mutually exclusive, for why the chr7-ins and chr19-ins Y alleles are not seen in the high PC1 benthic ecomorph. The first is that the PCA and admixture plots (Figure 3a, Supplementary Figures 4, 5) are consistent with an asymmetry of gene flow between the benthic and littoral ecomorphs, with the benthic ecomorph that is adapted to the cold, hypoxic environment at the bottom of the lake being genetically isolated with little if any gene flow from littorals into it, whereas there is gene flow from the benthics into littorals. This supports the cline of benthic admixture reflected in PC1 variation amongst the littorals. Second, even if there is hybridisation leading to low levels of gene flow into benthics, there are reasons to suggest it is sex-biased involving littoral females and benthic males. We never caught genetically benthic fish in the shallow depths where littorals breed, but we do see occasional genetic littorals in deep water. Benthic males appear to exclusively use the deep water mating territories that have been observed at the base of the crater wall, and we suggest that littoral males may be unable to compete successfully in this forbidding environment to which they are not adapted whereas littoral females may accept mating. In this scenario low frequency Y alleles from the littorals would not invade the benthics at an appreciable rate, and any that were present in the founders or entered through rare hybridization events could have been easily lost by drift.

In conclusion, our discovery that at least three different alleles control sex and segregate differentially within an isolated population of *A. calliptera* provides evidence that genetic sex determination in nature can be extremely fluid even at very small demographic scales. All of the alleles we identified involved structural genetic variants, with two of the three generated by transposable element insertions, highlighting a potentially important role for TEs in the rapidly evolving sex systems of African cichlids, similar to their role in adaptive variation in opsin regulation (Carleton *et al*. 2020). Our results also indicate that genetic background differences likely created by admixture can bring about antagonistic relationships among males carrying different Y alleles, providing an evolutionary context that may favour multifactorial sex systems. This has interesting implications for the incipient speciation between littoral and benthic Masoko ecomorphs in that alternative Y alleles circumvent negative genetic interactions brought about by admixture, allowing for sustained back-crossing that reduces the level of divergence. It is possible that this contributes to the low genome-wide F_ST_ (4%) between the ecomorphs, which also lack fixed genetic differences, although there are tens of islands of high F_ST_ divergence potentially associated with loci under differential selection (Malinsky *et al*., 2015). Admixture and relatively low divergence are hallmarks of the Malawi cichlid radiation, so it seems plausible that similar processes could exist or have existed elsewhere. The fact that we and other studies have found polygenic sex determination systems that differ markedly between closely related species and populations across the radiation supports this possibility.

## Methods

### Samples and sequencing

Fish were primarily collected by professional aquarium fish catching teams. Fish at a target depth range (determined by diver depth gauges) were chased into block nets by SCUBA divers and transferred to a holding drum, then brought to the surface, where they were euthanized with clove oil. The right pectoral fin of sampled individuals was then removed and stored in ethanol, and the remainder of the specimen pinned, photographed, labelled and preserved in ethanol for later morphological analysis. Standard lengths were measured using calipers. Females were distinguished from juvenile males among the smaller fish by visual inspection of the gonads after opening the abdominal cavity. Adult males were identified from secondary sexual traits of larger size, brighter colour and possession of elongate filaments on the pelvic, dorsal and anal fins (confirmed to be reliable by visual inspection of the gonads in a number of specimens from earlier collections).

DNA was extracted from preserved fin clips using Qiasymphony DNA tissue extraction kits or PureLink® Genomic DNA extraction kits and samples were sequenced on the Illumina HiSeq2000 as in Malinsky *et al*. (2015) or on the HiSeqX in three batches: 1) 118 “ILBCDS” samples collected in 2011 sequenced at 3.9-19.2x coverage (median 7.5x), 2) 194 “CMASS” samples collected in 2014-2016 sequenced to 4.3-9.0x coverage (median 5.7x), 3) 336 “cichl” samples collected in 2014-2016 and 2018 sequenced to 12.0-23.2x coverage (median 15.8x).

One sample that was initially part of the study was removed following conflicting data being detected during the analysis. Further testing with our PCR assay of both the original tissue sample obtained in the field, and a second sample from the supposed same ethanol-preserved, whole specimen, produced one male and one female genotype respectively, indicating a labeling error (Supplementary Figure 1c).

RNA was extracted from the gonads of two male and two female *A. calliptera* collected from the Itupi River in 2016. To ensure accurate quantification of transcripts, we used PolyA selection on one male and one female sample and RNA depletion on the other male and female sample. The gonad libraries were then sequenced using 75 bp paired-end reads on three lanes of the Illumina HiSeq 2500 (SBS kit v4). Adapter sequences and bases with Phred quality below 20 were removed from the ends of gonad RNAseq reads using Trim Galore 0.6.2 (https://www.bioinformatics.babraham.ac.uk/projects/trim_galore/) and read quality was checked using FastQC 0.11.8 (https://www.bioinformatics.babraham.ac.uk/projects/fastqc/). We also extracted RNA from the anal fins, eyes, gills and livers of 151 *A. calliptera* collected from Lake Masoko in 2015, 2016 and 2018 (Supplementary Table 1), which was stored in RNALater, using Direct-zol™ RNA MiniPrep Plus kits (Zymo, R2072) with an additional Chloroform step before loading the sample onto filtration columns. RNA samples were quantified with the Qubit™ RNA HS Assay Kit and quality assessed on the Agilent 4200 TapeStation. Libraries were prepared using Illumina mRNA sequencing kits with polyA enrichment and sequenced using 100 or 150 bp paired-end reads on three lanes of the Illumina HiSeq4000 and five S4 lanes of the Illumina NovaSeq. Adapter sequences and bases with Phred quality below 20 were removed from the ends of all resulting RNAseq reads using Trim Galore 0.6.4 and read quality was checked using FastQC 0.11.9.

### Variant discovery

Sequencing reads for all *A. calliptera* samples were mapped to a high-quality *A. calliptera* reference genome (fAstCal1.2, accession GCA_900246225.3) (Rhie *et al.*, 2021) using bwa-mem 0.7.17 (Li, 2013). We used GATK 3.8 (McKenna *et al*., 2010) to identify individual-level variation with the HaplotypeCaller program followed by joint genotype calling among all samples using GenotypeGVCFs (Poplin *et al*., 2017; Van der Auwera & O’Connor, 2020). Sites exhibiting any of the following indications of quality issues in the medium-coverage (~15x) “cichl” subset of 336 individuals were masked from all analyses: total sequencing depth across individuals more extreme than the genome-wide median total site depth (DP) +/−25%, fewer than 90% of individuals covered by at least eight reads, more than 10% of individuals with missing genotypes, root mean square mapping quality less than 40, an alternate allele assertion quality score below 30, a variant quality by depth score below three, excess heterozygosity (exact test p-value < 1e-4), biases between reference and alternate alleles in terms of strand (exact test p-value < 1e-6), base quality (z-score > 6), mapping quality (z-score > 6), and read position (z-score > 6). Sites spanning indels or having more than two alleles were also masked from analyses. Quality control for sites was carried out using the program vcfCleaner (https://github.com/tplinderoth/ngsQC/tree/master/vcfCleaner).

### Population genetic characterization

We used principal component analysis (PCA) based on genotype posterior probabilities at the quality-controlled SNPs to characterize the distribution of *A. calliptera* genetic variation throughout Lake Masoko. Specifically, we used ANGSD 0.929 (Korneliussen *et al*., 2014) to estimate minor allele frequencies from genotype likelihoods (-GL 1 model) calculated using reads with minimum base and map Phred qualities of at least 20. These minor allele frequency (MAF) estimates and genotype likelihoods were used to obtain genotype posterior probabilities for all individuals under a Hardy-Weinberg genotype prior. We used ngsCovar 1.0.2 (Fumagalli *et al*., 2014) to estimate the genetic covariance matrix among individuals based on their genotype posteriors at SNPs with MAF greater than 5%, which we decomposed in R 3.6.3 (R Core Team, 2020) with the eigen() function. In addition, we used the program ADMIXTURE 1.3.0 (Alexander *et al*., 2009) to infer the proportions of distinct genetic ancestry for individuals assuming two ancestral populations (K parameter).

### Genome-wide association tests for sex

We relaxed some quality filters to accept additional biallelic SNPs for statistical association testing by requiring that they have a minimum total depth across individuals of 2000x (lowered from 3500x), at least 90% of individuals covered by a minimum of four reads, and an exact test p-value for excess heterozygosity above 1e-20. All other quality criteria were kept the same. We queried all such SNPs across the genome with MAF of at least 5% for association with sex under the linear mixed model framework implemented in GEMMA 0.98.1 (Zhou & Stephens, 2012). Sex was treated as a binary response which we regressed against posterior mean genotypes calculated from the GATK genotype likelihoods using vcf2bimbam (https://github.com/tplinderoth/ngsQC/tree/master/vcfCleaner) under a Hardy-Weinberg genotype prior. We accounted for confounding effects of ancestry among individuals through incorporating a centered pairwise kinship matrix calculated using GEMMA as a random effect in the LMM. We identified significantly associated loci using the likelihood-ratio test p-values from GEMMA run in the LMM mode at a 5% significance level after a Bonferroni correction for the number of tested SNPs. In order to identify as many sex-associated loci as possible, we iteratively tested conditional subsets of individuals who did not carry alleles significantly associated with sex from previous iterations, that is, subsets of individuals whose sex was not accounted for by other candidates.

### Characterizing sex-determining variants throughout Lake Masoko and the Malawi radiation

We only used SNPs with GEMMA and so following the sex GWAS we checked for the presence of structural variants (SVs) that might have a stronger association with sex in 10 kb windows extending from the significantly associated SNPs. We extracted read mapping information directly from the BAM files to look for mapping signatures that would be consistent with structural variation, considering both read pair and depth information, using IGV 2.8.0 (Robinson *et a*l., 2011). We initially screened at least five males and five females for structural variation in IGV and then used a custom perl script to call SVs if at least 5% of read pairs among all individuals within 480 bp of any putative SV positions had mates which mapped to a different chromosome. We assembled the anomalously mapped read pairs across all individuals for each SV that we called using MEGAHIT 1.2.9 (Li *et al*., 2016) and performed a blastn (Altschul *et al*., 1990; Camacho *et al*., 2009) search of the resulting contigs against fAstCal1.2. This approach led to the discovery of the putative sex-determining insertions on chromosomes 7 and 19, which blasted with at least 90% identity across their full length to multiple places across the genome. We used repeatModeler2 2.0.2 (Flynn *et al*., 2020) with default options but including the-LTRStruct option to identify transposable element sequences in the fAstCal1.2 genome. Then we compared the SV contigs to these transposable element sequences to further characterize the insertions. The chr19-ins allele matched a 700 bp transposable element (blastn evalue = 0, 97% identity, 99% coverage) identified by repeatModeler2 as belonging to an LTR/Unknown family. The two partial contigs of the chromosome 7 insertion matched with 94% identity (631/673 bp with 35/673 bp (5%) gaps) and 97% (496/509 bp with 11/509 bp (2%) gaps) to either end of a 3,947 bp unknown transposable element.

In order to characterize the presence or absence of the chromosome 7 and 19 insertions, we mapped sequencing reads from all Masoko *A. calliptera* to the assembled insertion sequences including 1 kb of upstream and downstream flanking sequence using BWA. We considered any reads mapping within the flanking regions and which spanned the insertion as reference allele reads (with respect to fAstCal1.2) and any reads which mapped within the insertion by a minimum of three bp as alternate allele reads. An individual’s genotype was called heterozygous (0/1) if they possessed reads from both alleles that were each at a minimum frequency of 10%, otherwise, with more than 90% of either the reference or insertion reads, individuals were called as homozygous for the reference allele (0/0) or homozygous for the insertion allele (1/1), respectively. We also genotyped fish based on the copy number of the duplicated *gsdf*-containing locus which spans positions 18,079,155 to 18,100,834 of chromosome 7 in the fAstCal1.2 reference. For each individual, we translated their average sequencing depth across this region relative to their average sequencing depth from 38,320 bp flanking sequence (19,154 bp upstream and 19,166 bp downstream of the duplication breakpoints) into copy number in increments of 0.5x: Relative coverage of 1.25 or lower was recorded as a non-duplicated *gsdf* region, (1.25,1.75] as three *gsdf* copies, (1.75, 2.25] as four copies, and so on. Individuals with three and four copies of the *gsdf* locus were called heterozygous and homozygous for the duplication respectively. Though it is possible for a four-copy individual to have one chromosome with three *gsdf* copies this would necessitate another duplication and so is less parsimonious than the assumption that they are homozygous for a chromosome with two copies.

We also developed a PCR assay for the *gsdf* duplication (Supplementary Table 7), which we used to confirm its presence in a subset of *A. calliptera* and *Maylandia zebra*. Genomic DNA was extracted from fin clips using PureLink Genomic DNA Mini Kits (ThermoFisher Scientific, K182001) following the manufacturer’s protocols and eluted in 30-60 μL elution buffer. We carried out PCRs in 20 μL reaction volumes consisting of 1X Platinum™ II PCR Buffer, 0.2 mM of each dNTP (ThermoFisher Scientific, R0192), 0.2 μM of each primer (Merck Life Science, desalted), less than 500 ng template DNA (1 μL genomic DNA at ~1-5 ng/μL), 0.04 U/μL Platinum™ II Taq Hot-Start DNA Polymerase (ThermoFisher Scientific, No 14966001) and nuclease-free water. We amplified the DNA using the following thermal profile: 94°C for two minutes followed by 30-35 cycles of 94°C for 15 seconds, 60°C for 15 seconds, 68°C for 15 seconds, and a final 68°C extension for five minutes. The PCR products were separated using electrophoresis run at 100 volts for 30 minutes on a 2% agarose gel.

We genotyped 1,552 additional individuals from all seven of the Lake Malawi radiation clades (*A. calliptera*, Mbuna, Benthic, Deep, Utaka, *Diplotaxodon*, and *Rhamphochromis*; see Malinsky *et al.* 2018) for the *gsdf* duplication as well as the chromosome 7 and 19 insertions in the same way as for Masoko *A. calliptera* described above. This set of Malawi radiation individuals represents 255 species (some are not formally established but recognized as distinct taxa) from 47 genera, including *A. calliptera* from locations other than Lake Masoko. In order to characterize how the *gsdf* duplication is acquired and lost as lineages diversify we mapped its presence at different copy number in males and females to the species tree for four Mbuna species from the *Maylandia* genus: *M. zebra*, *M. callainos*, *M. emmiltos*, and *M. fainzilberi*. We generated the species tree using 12,133,030 genome-wide segregating sites among the four *Maylandia* species identified using GATK 3.8 in the same manner as for Masoko *A. calliptera*. These SNPs passed quality controls addressing abnormally low and high sequencing coverage and low mapping quality for the ingroup samples as well as for samples from the distantly-related species *Rhamphochromis longiceps,* which served as an outgroup. We used ngsDist 1.0.8 (Vieira *et al*., 2016) to calculate a pairwise genetic distance matrix based on genotype likelihoods for all of the ingroup and outgroup samples, as well as to bootstrap sites in order to generate 100 additional bootstrap distance matrices. For this *Maylandia* species tree, we used fastME 2.1.6.1 (Lefort *et al*., 2015) to infer neighbor-joining trees from the genetic distance matrices using the BIONJ algorithm with SPR tree topology improvement. RAxML-NG 1.0.1 (Kozlov *et al*., 2019) was used to determine the bootstrap support for the genome-wide tree.

### B chromosome assay

In addition to autosomal sex loci, B chromosomes, which are supernumerary chromosomes not required for organismal function and variably present across taxa and individuals, have been implicated as sex modifiers in Lake Malawi cichlids (Clark *et al*., 2017). Accordingly, we assayed for the presence of B chromosomes among Masoko *A. calliptera* to discern whether they may influence sex. B chromosome material initially derives from autosomes, so their presence can be detected through inflated read coverage in homologous regions of the reference genome where B reads mismap. Accordingly, we assayed for B chromosomes based on inflated coverage at regions containing sequence known to exist on B chromosomes from Lake Malawi cichlids (Clark *et al*., 2018). Regions identified as core B block sequence according to Clark *et al*. (2018) were translated into fAstCal1.2 coordinates and the mean coverage across each of these segments for each Masoko *A. calliptera* individual was calculated directly from the BAM files. We used a minimum coverage ratio for the core B region compared to the genome-wide average of 2x to call B positive individuals. None of the Lake Masoko *A. calliptera* passed this threshold although this process did identify individuals carrying B chromosomes from other species.

### Expression of sex-associated genes

We mapped the quality-controlled liver, eye, gill, and anal fin RNAseq reads to the fAstCal1.2 genome with STAR 2.7.3a (Dobin & Gingeras, 2015) and counted reads derived from sex-associated genes with featureCounts 2.0.1 (Liao *et al*., 2014). These read counts were normalized to counts per million (CPM) reads using edgeR 3.30.3 (Robinson et al., 2010). We mapped the quality-controlled gonad reads to the fAstCal1.2 reference using bwa-mem and counted reads derived from *gsdf* exons using SAMtools 1.9 (Li *et al*., 2009) and ngsAssociation 0.2.4 (https://github.com/tplinderoth/ngsAssociation) summarize, which were also normalized to CPM.

### Relationship between Y alleles and body size

Genetic PC1 was used as a proxy for the degree of admixture since this component clearly separates fish based on their degree of benthic ancestry. Based on distinct clustering in the genome-wide PCA plot, fish with PC1 > 0.04 were classified as genetically benthic and those with PC1 < 0.04 as genetically littoral. We further classified fish with the lowest amounts of benthic ancestry as “low PC1” (PC1 < −0.02), those with more equal amounts of littoral and benthic ancestry as “middle PC1” (PC1 range −0.02 to 0.04), and the clear benthic cluster as “high PC1” (PC1 > 0.04). The three Y alleles segregate in the littoral group only, which is composed of low and middle PC1 fish, yielding six possible Y and PC1 combinations when excluding the 0.7% of males that carry more than one type of Y. For all analyses related to fish size we considered only males that were heterozygous for their Y allele (except when we compared the length of *gsdf*-dup homozygotes to *gsdf*-dup heterozygotes). We tested the hypothesis that littoral Lake Masoko *A. calliptera* males with different ancestry backgrounds and Y allele combinations differ in standard length using pairwise two-tailed t-tests in R.

We investigated whether the size of littoral males is influenced by interactions between Y allele and ancestry regime by fitting linear models of standard length as a function of Y allele and PC1 class in R using glm(). We tested whether the interaction provides a significantly better fit with the anova() F-test by comparing the residual sums of squares between a model with only main effects to a model with main effects and an interaction between Y allele type and PC1 class. We also introduced a depth class variable into our models to investigate whether the depth at which fish were caught plays a role in explaining their length. Depths less than five metres were considered “shallow”, depths ranging from 5-20 metres were “intermediate”, and depths more than 20 metres were “deep”. As before, we compared the fit of a saturated model including the three-way interaction between Y allele, PC1 class, and depth band to the same model but without the three-way interaction using analysis of variance to determine if the joint interaction between all variables provides a significant amount of additional power for predicting fish length.

Since the size of male fish is likely to influence fitness, we used log-linear models to look at whether the same factors affecting length could predict the frequency of males. Specifically, we fit models using glm() in R with family=’poisson’ for the frequency of males based on Y allele, PC1 class, and depth band. We assessed whether the frequency of males belonging to categories based on these three variables are independent of one another, and if not, what interactions were involved by performing an analysis of variance on nested pairs of models. We tested whether the differences in the residual deviance between the models being compared were significant using χ^2^ tests. This enabled us to find the simplest model that predicts male frequencies statistically as well as the saturated model that includes all main effects and their possible interactions. The significance of terms within the context of a particular model for which they were fit was determined using a Wald test of the null hypothesis that a term’s effect is equal to zero.

### Assessment of linkage disequilibrium around sex loci

We calculated LD in terms of r^2^ between each of the most highly sex-associated GWAS SNPs and their surrounding SNPs using PLINK 1.9 (Purcell, 2014; Purcell *et al*., 2007). We observed high LD, r^2^ > 0.5, between the strongest GWAS SNPs tagging chr19-ins and chr7-ins and far-ranging surrounding SNPs, which we visualized using plot_zoom (https://github.com/hmunby/plot_zoom). In order to determine how unusual these long stretches of high LD were, we compared the variance in the pairwise physical distance between the top GWAS SNPs and all SNPs within one megabase and r^2^ > 0.5 to an expected distribution. The background distributions were generated by randomly sampling 5,000 focal SNPs from across the genome having the same alternate allele frequencies as each of the top GWAS SNPs. For each sampled SNP, we calculated the variance among pairwise distances with other SNPs in the same way as we had done for the GWAS SNPs.

## Acknowledgments

We are grateful to African collaborators who assisted in sample collection, particularly the staff of the Tanzanian Fisheries Research Institute, as well as Alan Hudson. We thank the sequencing core staff at the Wellcome Sanger Institute. This work was supported by the Wellcome Trust (WT207492 and WT206194). Additional support was to MJG & GFT Leverhulme Trust - Royal Society Africa Awards (AA100023 and AA130107); to MJG Leverhulme Trust award (RF-2014-686); to GFT Leverhulme Trust award (RPG-2014-214); to EAM Wellcome Trust Senior Investigator award (104640/Z/14/Z and 219475/Z/19/Z) and CRUK award (C13474/A27826). GV thanks Wolfson College, University of Cambridge and the Genetics Society, London for financial support.

---

**Figure S1:**
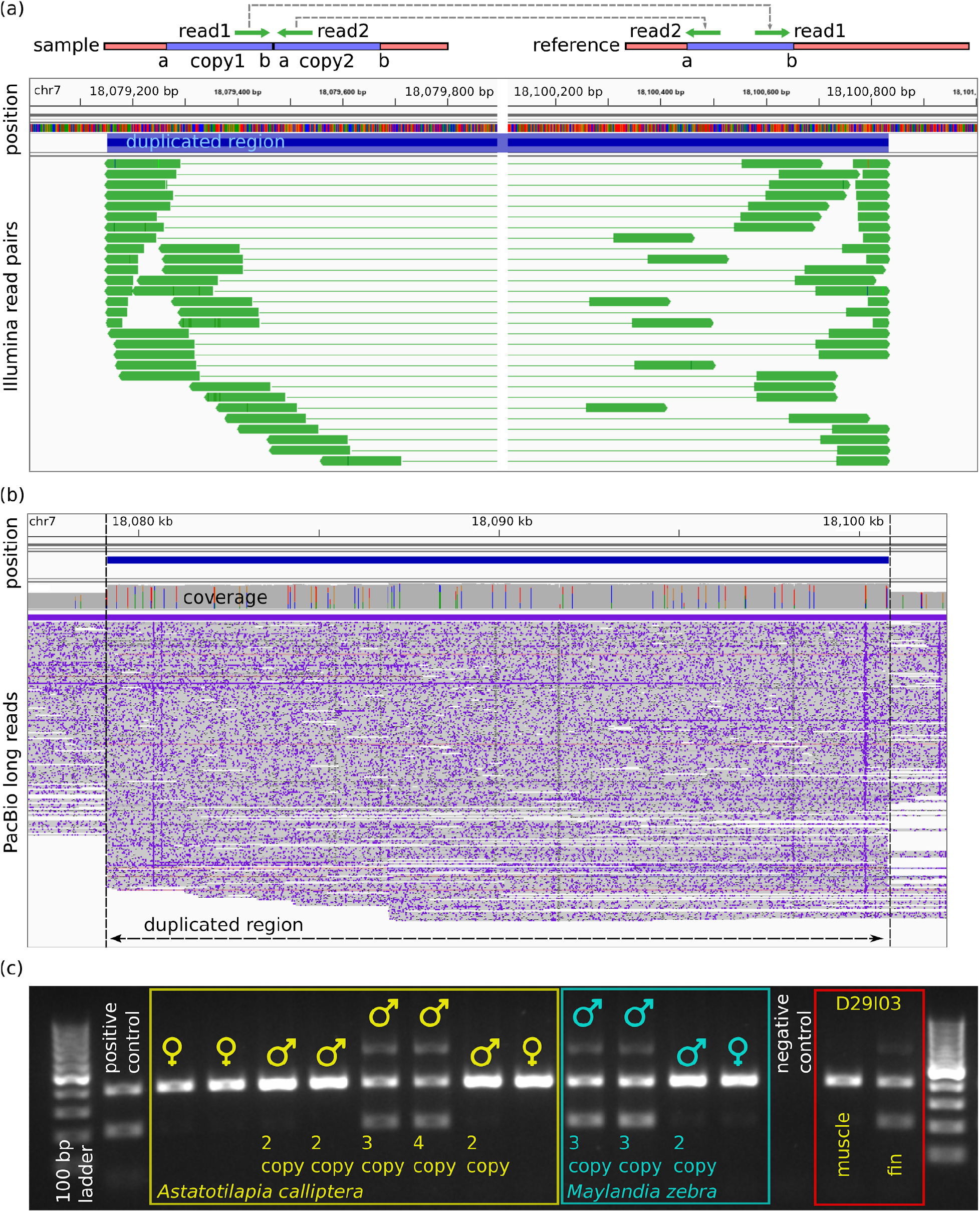
Characterization of the *gsdf* duplication. **(a)** Short Illumina reads from four Masoko male *A. calliptera* called homozygous for the *gsdf* duplication based on relative sequencing depth that is approximately 2x higher than in ~38 kb of non-duplicated flanking sequence. The mapping orientation of all read pairs to the fAstCal1.2 reference is consistent with a tandem duplication as shown in the schematic at the top. **(b)** PacBio reads from a male *Tropheops ‘mauve’* mapped to the fAstCal1.2 reference. The sharp break in the alignment of some of the reads at the edges of the *gsdf* duplication (blue horizontal bar) in conjunction with elevated coverage signals that this individual is heterozygous for the same *gsdf* duplication identified in Masoko *A. calliptera*. **(c)** Agarose gel image of PCR products from primers designed to assay for the presence of the *gsdf* duplication. Based on this assay, individuals positive for the *gsdf* duplication yield three distinct bands, whereas those negative for the duplication produce a single band. The assay was used to confirm the presence of the duplication in two male *Maylandia zebra* samples that were putative heterozygotes for *gsdf*-dup based on sequencing depth. Two separate tissues for Masoko *A. calliptera* sample D29I03 produced different genotypes based on this PCR assay indicating a sampling error and resulted in this individual being omitted from all analyses.

**Figure S2:**
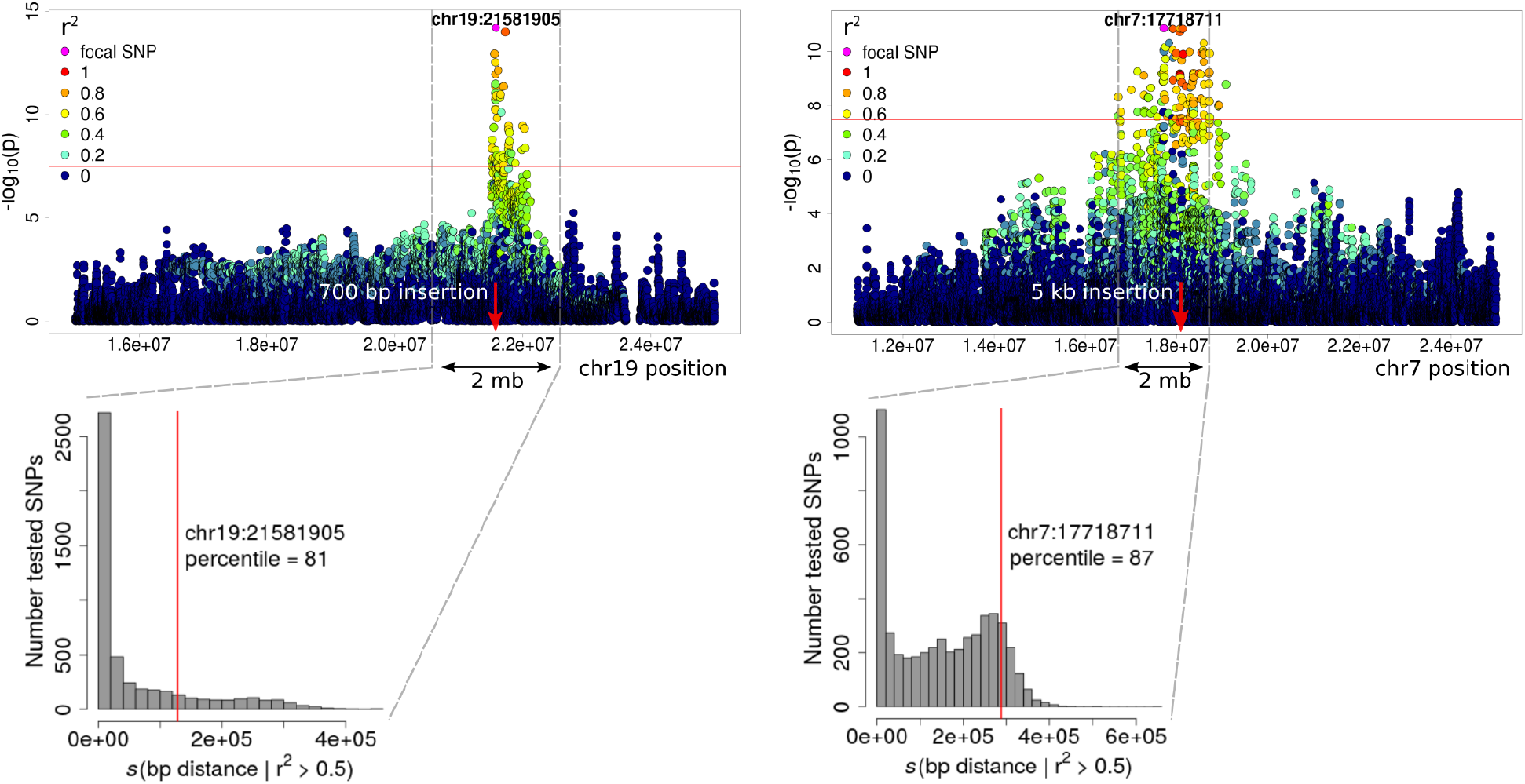
Elevated linkage disequilibrium around the chr19-ins and chr7-ins loci. The top Manhattan plots are a regional view of the p-values for the likelihood ratio test from the GWAS for sex used to identify SNPs tagging chr19-ins (left) and chr7-ins (right). The positions of the insertions are denoted with red arrows. Elevated linkage disequilibrium (LD) between the SNP with the highest sex association in each GWAS and other surrounding SNPs extends far along the respective chromosomes. This causes the variance in the pairwise physical distance among SNPs in high LD (r^2^ > 0.5) with the top GWAS SNPs to be higher than typically expected throughout the genome, consistent with recent positive selection. The histograms show where this variance for the top GWAS SNPs fall along the expected distributions for Masoko *A. calliptera*, which were generated by randomly sampling 5,000 SNPs across the genome with the same alternate allele frequencies as the GWAS SNPs. The variance among the pairwise distances between each sampled SNP and their surrounding high-LD SNPs were calculated in the same manner as for the GWAS SNPs.

**Figure S3:**
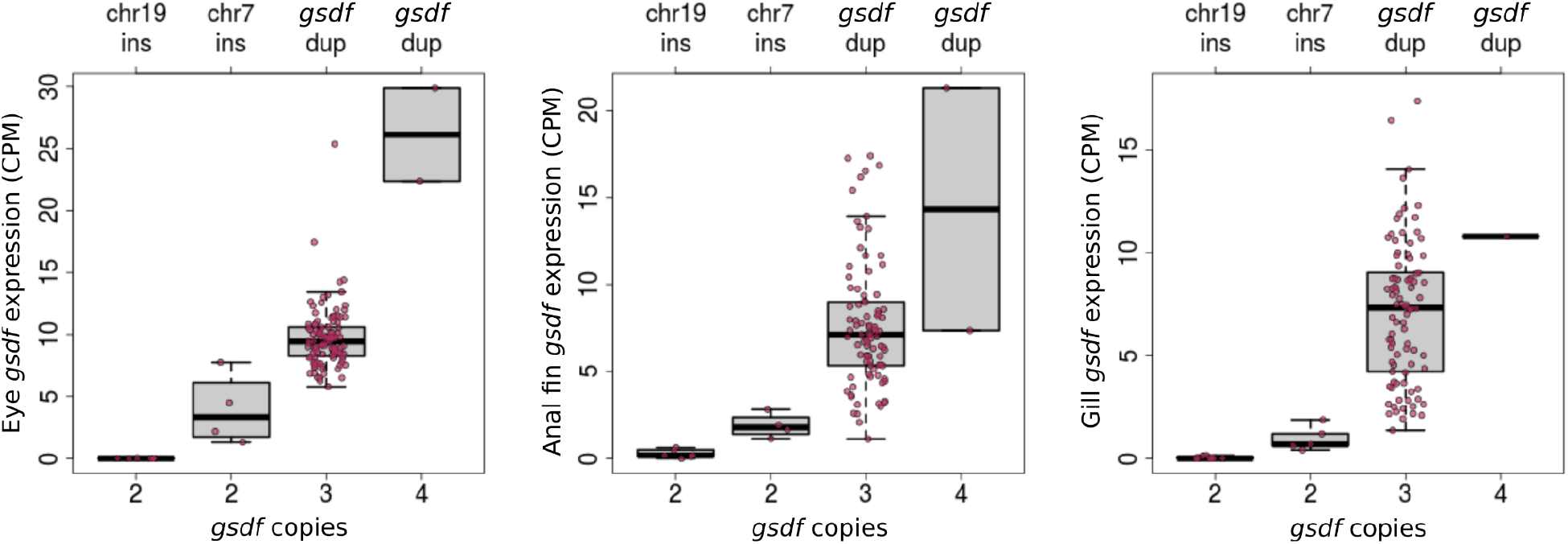
Expression of *gsdf* in somatic tissues for males with different Y alleles. The *gsdf*-dup and chr7-ins alleles are defined by a tandem duplication of the *gsdf* gene and an insertion directly upstream of *gsdf*, respectively. Levels of *gsdf* expression in eye, anal fin, and gill tissues from Masoko male *A. calliptera* demonstrate that males carrying putative Y alleles generated through mutations involving *gsdf* express this gene more than other males.

**Figure S4:**
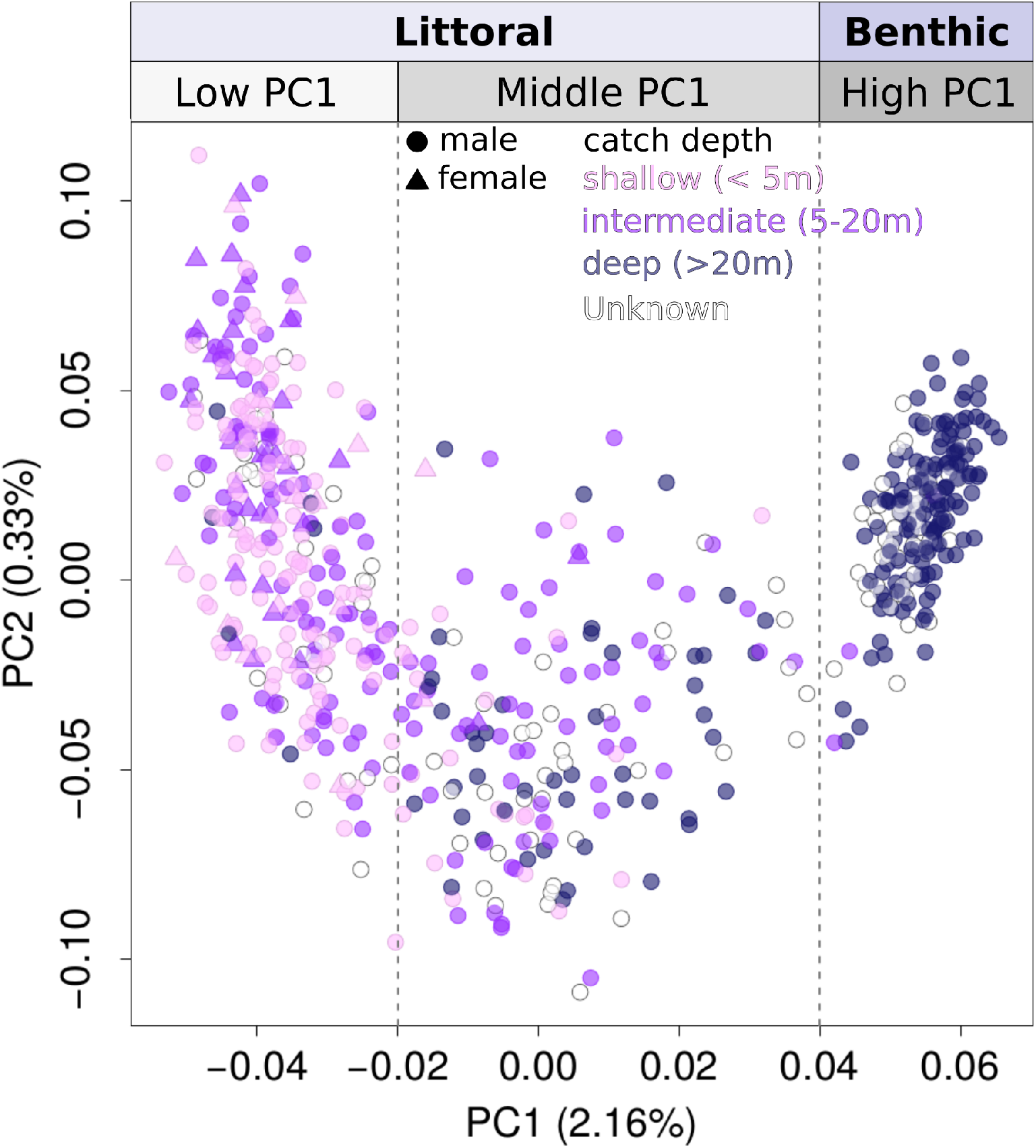
Relationship between genetic variation and catch depth. Lake Masoko *A. calliptera* distributed along the first two components of a principal component analysis of genome-wide variation reveals strong philopatry of high PC1 fish for deep depths. This coincides with nearly all high PC1 individuals conforming to the benthic ecomorph. In contrast, fish below PC1 values of 0.04 are almost all of the littoral ecomorph and exhibit far less constrained habitat preference. Among littoral fish (PC1 < 0.04), the most admixed individuals in the middle of PC1 (−0.02 to 0.04) regularly occupy all depth bands, while low PC1 littorals (PC1 < −0.02) remain mostly at depths above 20 metres, though occasionally they are found deep.

**Figure S5:**
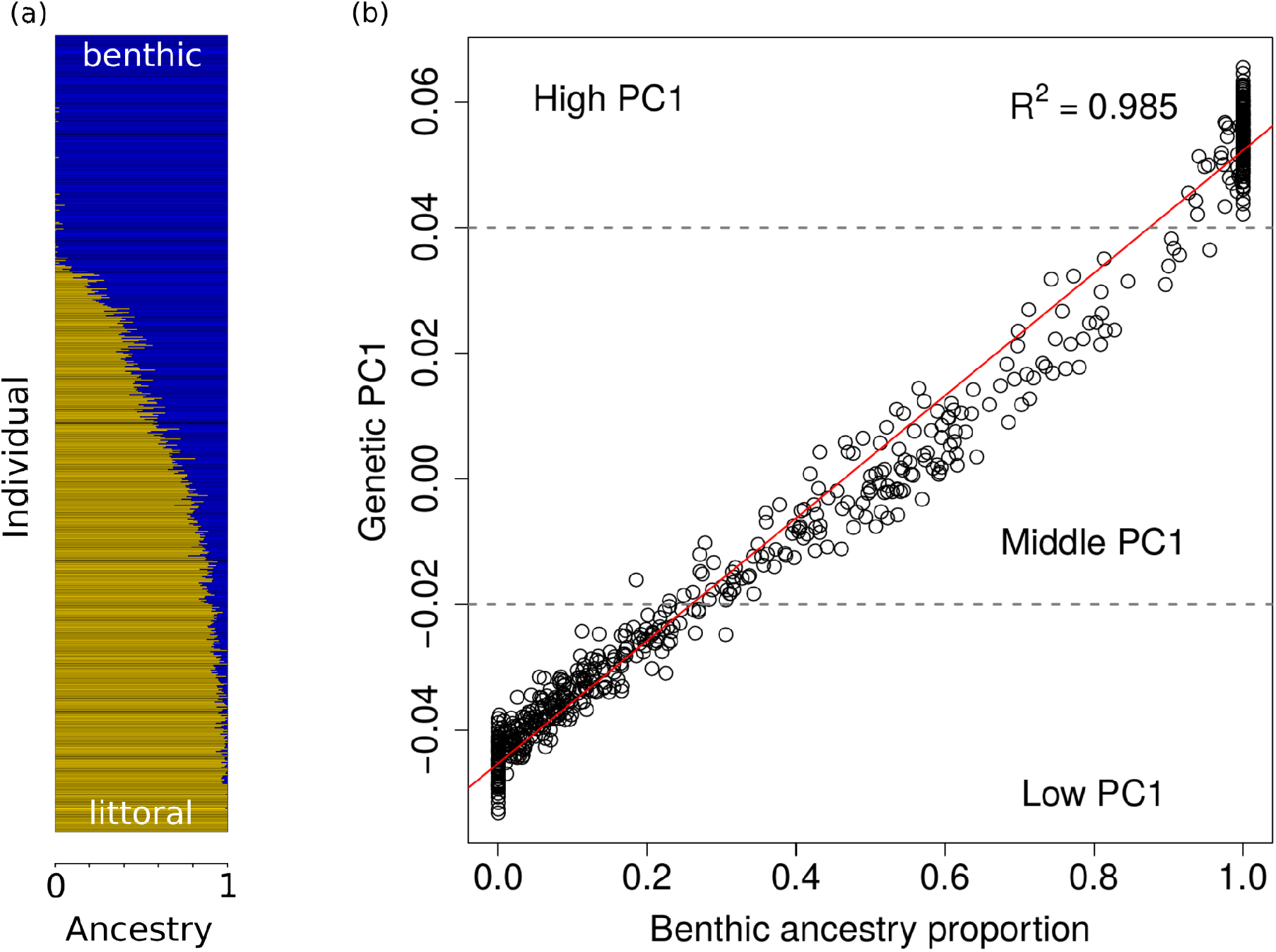
Ancestry characterization of Masoko *A. calliptera*. **(a)** Genome-wide ancestry proportions for individuals inferred using the program ADMIXTURE and ordered by their genetic PC1 rank shows the genetic distinctiveness of the benthic (high PC1) subgroup, a subset of littorals having low amounts of benthic ancestry (low PC1), and a highly admixed group (middle PC1). **(b)** The genetic PC1 scores of Lake Masoko individuals regressed against their proportion of benthic ancestry shows that PC1 almost perfectly describes the genetic structure of the Lake Masoko population in terms of the continuum between genetically benthic and littoral ancestries. The fitted linear regression line is shown in red and the low, middle, and high PC1 classification cutoffs are depicted with dashed grey lines.

**Figure S6:**
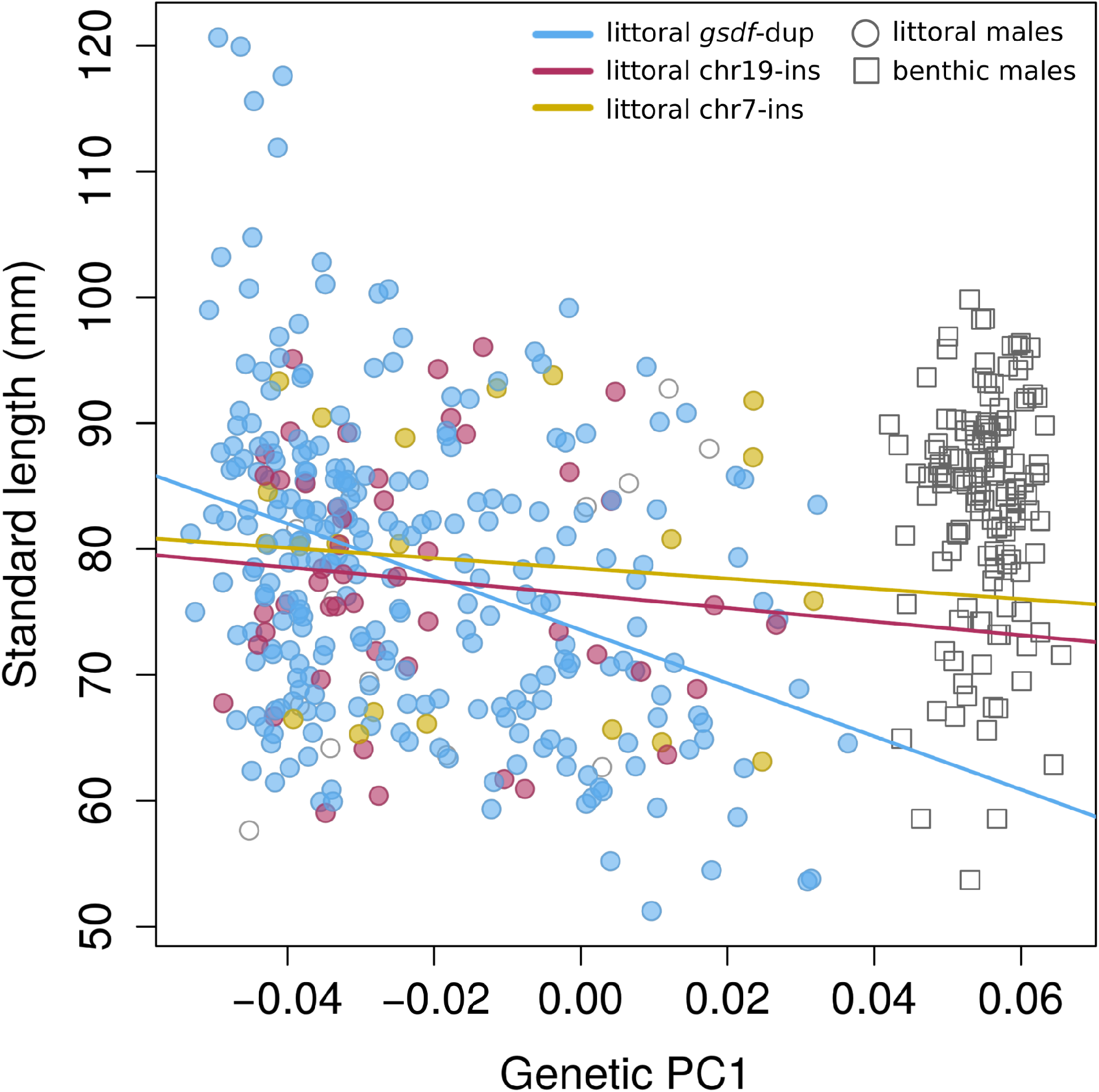
Interaction between genetic background and Y allele in predicting male size. The standard lengths of male *A. calliptera* from Lake Masoko plotted against their position along PC1 of the principal component analysis of genome-wide variation shows a negative trend in the length among genetically littoral (PC1 < 0.04) males (circles) with increasing PC1 value. Linear regression models of length predicted by PC1 were fitted separately for littoral males heterozygous for either *gsdf*-dup, chr19-ins, or chr7-ins corresponding to the colours blue, red, and yellow, respectively. Littoral males carrying more than one Y allele, homozygous for Y alleles, or which did not have an identified Y, are represented by uncoloured circles and were excluded from the regressions. Genetically benthic males, defined as fish with PC1 > 0.04, are plotted for comparative purposes as squares without any indication of their Y genotype. The distinctly more negative slope of the regression line fit to *gsdf*-dup males compared to chr19-ins and chr7-ins males shows that length is predicted to decrease much more drastically with more benthic admixture among *gsdf*-dup males. This difference is so great that males using *gsdf*-dup are predicted to switch from being longer than males using other Y alleles to actually being shorter above PC1 values of −0.02.

**Figure S7:**
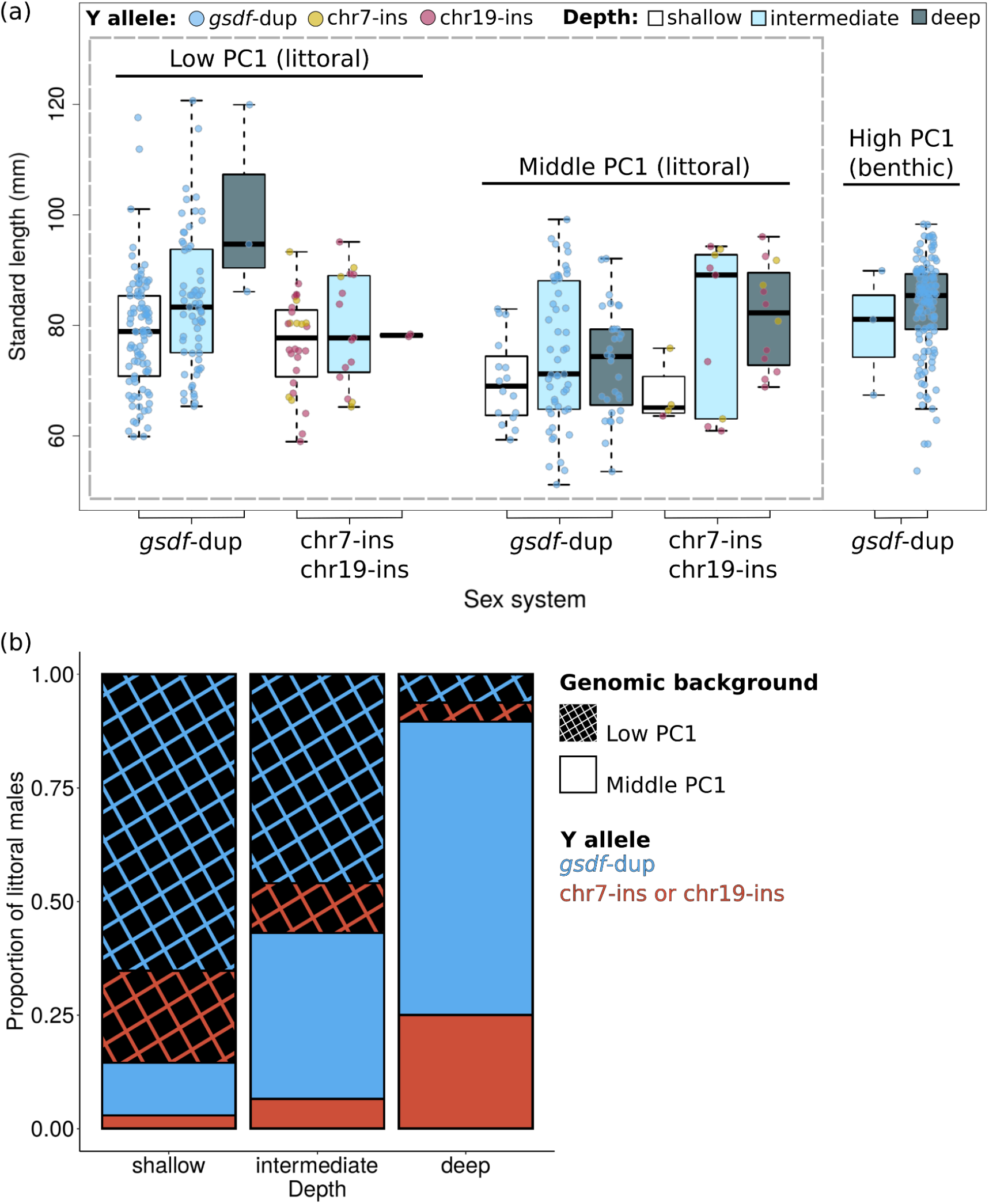
Male sizes and frequencies according to Y allele, genetic PC1, and catch depth. **(a)** Standard length comparisons across different PC1 genetic backgrounds and catch depths of Lake Masoko *A. calliptera* males heterozygous for only one of the Y alleles shows an interaction between Y allele, catch depth, and PC1 background in predicting size. Among the genetically littoral males (within the dashed grey box) those carrying *gsdf*-dup are smaller on middle PC1 versus low PC1 backgrounds regardless of what depth they are found at. In contrast, among males using the other Y alleles only middle PC1 males found in shallow waters are smaller than the low PC1 males, while at deeper depths their size remains constant across genetic backgrounds and may even show a subtle tendency to be larger with middle PC1 benthic ancestry. **(b)** A comparison of the proportion of littoral males characterized by different genetic PC1 backgrounds and Y alleles at different catch depths shows that the proportion of males with middle PC1 ancestry increases with depth. However, within PC1 backgrounds, the fraction of males using the different Y alleles remains relatively stable across depths. Overall, *gsdf*-dup males dominate at all depths.

**Tables S1 to S7** can be found in the attached Excel file: supplementary_tables_differential_use_of_multiple_genetic_sex_determination_systems_in_div ergent_ecomorphs_of_an_African_crater_lake_cichlid.xls. For convenience the table legends are given below, and we also copy below the contents of tables S3 and S7, which are short.

**Table S1: Lake Masoko *Astatotilapia calliptera* samples** Genetic, phenotypic, collection, and data availability information for all Lake Masoko *A. calliptera* samples. RNAseq expression levels for *gsdf* are reported in counts per million reads mapped (CPM). Sample accessions are provided for whole-genome (WGS) and RNAseq sequence data deposited into the European Nucleotide Archive. Missing values are coded as “NA”.

**Table S2: GWAS multilocus sex determination genotype frequencies** Counts of Masoko *A. calliptera* individuals, stratified by sex and PC1 genetic background, for all observed combinations of *gsdf* copy number and genotypes at the most strongly associated SNPs in the serial GWAS for sex. 0 = reference allele, 1 = insertion allele, ./. = missing genotype.

**Table S3:** Average sizes of Masoko males. The mean standard length of Masoko *A. calliptera* males heterozygous for one type of Y allele stratified by PC1 genetic background and catch depth.

| Lake-wide mean length (mm) |  |  |
| --- | --- | --- |
| Y allele | Low PC1 | Middle PC1 |
| *gsdf*-dup | 81.34 | 73.55 |
| chr7-ins or chr19-ins | 77.68 | 78.73 |
| **Shallow (< 5 m) mean length (mm)** |  |  |
| Y allele | Low PC1 | Middle PC1 |
| *gsdf*-dup | 78.55 | 69.91 |
| chr7-ins or chr19-ins | 76.67 | 67.46 |
| **Intermediate (5-20 m) mean length (mm)** |  |  |
| Y allele | Low PC1 | Middle PC1 |
| *gsdf*-dup | 84.41 | 74.87 |
| chr7-ins or chr19-ins | 79.50 | 79.96 |
| **Deep (> 20 m) mean length (mm)** |  |  |
| Y allele | Low PC1 | Middle PC1 |
| *gsdf*-dup | 100.26 | 73.33 |
| chr7-ins or chr19-ins | 78.22 | 81.56 |

**Table S4: Littoral male frequencies according to genetic type and catch depth** Counts of Lake Masoko *A. calliptera* littoral males heterozygous for one type of Y allele stratified by genetic PC1 background and depth at which they were caught.

**Table S5: Sex loci genotype calls for Lake Malawi cichlid radiation species** The number of *gsdf* copies and genotype (GT) calls for chr19-ins and chr7-ins (0 = reference allele, 1 = insertion allele, ./. = missing genotype) for individuals of different species belonging to the Lake Malawi haplochromine cichlid radiation. The AC values indicate the number of “<reference allele>,<insertion allele>” sequencing reads observed for an individual. Missing values are coded as “NA”.

**Table S6: Frequency of chr7-ins in non-*calliptera* species from the Lake Malawi haplochromine radiation** Counts of individuals from all species apart from *Astatotilapia calliptera* in which chr7-ins was found, stratified by *gsdf* copy number and chr7-ins genotype. Multilocus genotype calls are defined as <number of *gsdf* copies>/<number of chr7-ins alleles>: for example, “3/1” denotes an individual possessing three *gsdf* copies and who is heterozygous for the insertion allele at the chr7-ins locus. Genotype class cells with non-zero counts are highlighted for readability.

**Table S7:**
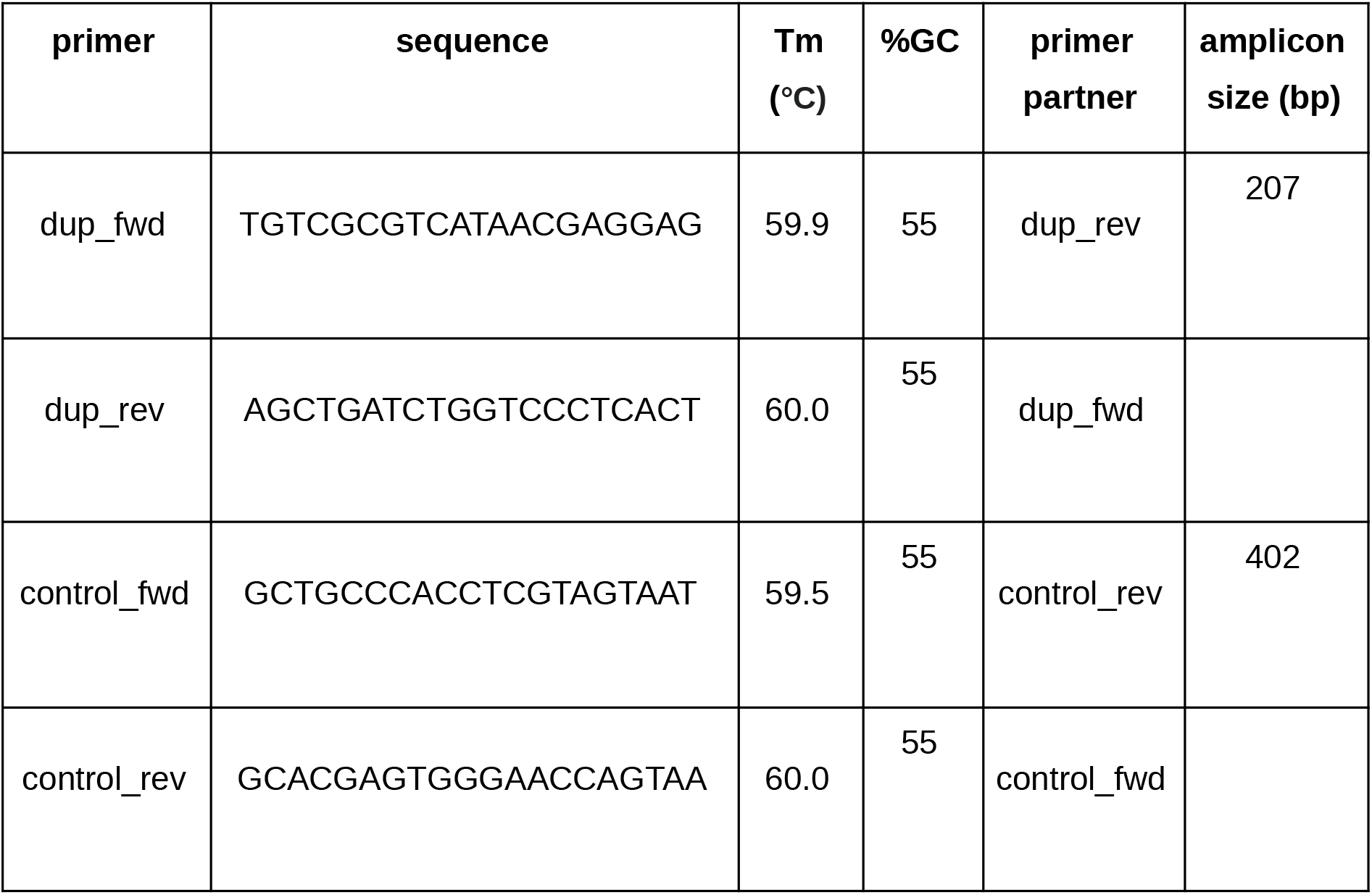
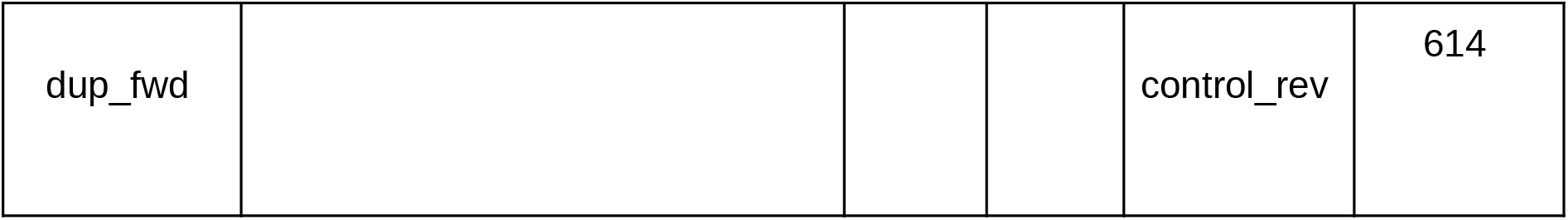
PCR primers for the detection of *gsdf*-dup. All samples should undergo amplification for the 402 bp control fragment, whereas only samples positive for the g*sdf* duplication should show equally strong amplification for the 207 bp fragment (and an additional 614 bp fragment which is not present when each primer pair is run in individual reactions).

